# Organization and regulation of nuclear condensates by gene activity

**DOI:** 10.1101/2022.09.19.508534

**Authors:** Halima H. Schede, Pradeep Natarajan, Arup K. Chakraborty, Krishna Shrinivas

**Affiliations:** School of Life Sciences, École Polytechnique Fédéral Lausanne, CH-1015, Lausanne, Switzerland; Department of Chemical Engineering, Massachusetts Institute of Technology, Cambridge, MA, 02139, USA; Institute for Medical Engineering and Science, Massachusetts Institute of Technology, Cambridge, MA, 02139, USA; Department of Physics, Massachusetts Institute of Technology, Cambridge, MA, 02139, USA; Department of Chemistry, Massachusetts Institute of Technology, Cambridge, MA, 02139, USA; NSF-Simons Center for Mathematical & Statistical Analysis of Biology, Harvard University, Cambridge, MA 02138, USA

**Author notes:** co-first authors (Halima H. Schede, Pradeep Natarajan).

## Abstract

Condensation by phase separation has recently emerged as a mechanism underlying many nuclear compartments essential for cellular functions. Nuclear condensates enrich nucleic acids and proteins, localize to specific genomic regions, and often promote gene expression. How diverse properties characteristic of nuclear condensates are shaped by genome organization and activity is poorly understood. Here, we develop a physics-based model to interrogate this interplay between condensation, active transcription, and genome organization. We show that spatial clustering of active genes enables precise localization and *de novo* nucleation of condensates. We find that strong clustering and activity drives aspherical condensate morphologies. Condensates flow towards distant gene clusters and competition between multiple clusters lead to stretched morphologies and activity-dependent repositioning. Overall, our model predicts and recapitulates morphological and dynamical features of diverse nuclear condensates and offers a unified mechanistic framework to study the interplay between non-equilibrium processes, genome structure, and multicomponent condensates in cell biology.

## Introduction

The cellular milieu is organized into dozens of membraneless compartments or biomolecular condensates, many of which form through phase separation^1–3^.Condensates concentrate multiple yet specific biomolecules through a network of multivalent and dynamic interactions^3–5^. Further, condensates exhibit a wide variety of physical and material properties and are actively regulated across the cell cycle^1,2,6^. In the crowded cellular environment, distinct condensates are coupled to non-equilibrium processes such as ATP-dependent chemical fluxes and mechanical remodeling that modulate their emergent properties^7,8^. This is particularly evident in the nucleus, where condensates interact with and are regulated by the genome, a large polymeric assembly of proteins, DNA, and RNA. The genome is intrinsically multi-scale and exhibits many layers of organization and regulation - from nanoscale nucleosomal clutches and microloops, to larger micron-scale compartments domains of active and inactive genes, and nucleus-scale territories for individual chromosomes^9–12^. Across these scales, genome organization is both modulated by and directly modulates active nuclear condensates. Examples include condensates that broadly promote gene expression such as the nucleolus, Histone locus body, nuclear speckles, and transcription-associated condensates, many first observed over a century ago ^13–21^. While many studies have emerged in the past decade to probe genome organization^9–12^ and condensates^1–3^ individually, our understanding of the coupling between genome organization, ATP-dependent transcription, and nuclear condensates remains nascent. More generally, how this interplay between active processes, multicomponent interactions, and heterogeneous environments dictate condensate properties is poorly understood.

Interactions between RNA and proteins are central in driving the condensation of nuclear bodies^22–25^, which in turn promote active transcription at specific genomic loci. Examples include: (1) Histone locus bodies (HLBs) enriched in transcriptional and regulatory proteins well as multiple genic, enhancer, and small nuclear RNAs that form around the histone gene cluster^26^ (2) Transcription-associated condensates which concentrate the transcriptional apparatus as well as noncoding and mRNAs, and preferentially localize to regulatory DNA elements called super-enhancers^27,28^ (3) Nuclear speckles which are enriched in the splicing apparatus as well as poly-adenylated mRNAs^29,30^ and interfacially localize particular subsets of active genes^29,31,32^. Many nuclear condensates assemble in a manner dependent on active transcription^23,33–35^ and exhibit stereotypic localization in the nucleus^24,36–40^. Further, emerging evidence indicates that low and specific levels of non-coding RNA may contribute to the formation of particular genomic or nuclear compartments^24^. How specific gene compartments modulate localization or nucleation of condensates by RNA is not well understood.

Unlike simple uniform liquids, nuclear condensates exhibit a wide gamut of transcription-dependent morphologies such as vacuoles^41,42^, aspherical shapes^30,39^,and layered organization of molecules^43–45^ that have been documented over many decades yet how these morphologies arise remain poorly understood. Nuclear condensates also exhibit unusual dynamics including bursts of directed motion ^46–49^, and inappropriate or aberrant morphologies of nuclear condensates often reflect pathological cell states^39,50,51^. Overall, nuclear condensates are highly variable transcription-dependent compartments with diverse morphologies, dynamics, and localization. Despite their central role in genome regulation, we do not have a unified mechanistic framework to study the emergent properties of nuclear condensates, in large part due to a lack of physically-grounded models of the underlying biology.

In this paper, we build a physically motivated *in silico* model to explore how spatial clustering of gene activity modulates condensate properties and thus transcription and nuclear organization. Through simulations, we first identify that compartmentation or clustering of active genes is sufficient to spatially localize nuclear condensates. At low rates of transcription, we find that clustered genes, through a positive feedback mechanism between local RNA synthesis and resultant protein recruitment, drives *de novo* nucleation of condensates. At higher rates of transcription from compartmentalized genes, we find that condensates adopt a range of asymmetric and non-equilibrium steady-state morphologies. When condensates are not proximal to gene compartments, we show that condensates can flow towards distant sites, driven by RNA gradients, and subsequently rationalize the limits of this directed motion through simple physical calculations. Finally, we show that relative clustering, activity, or separation between multiple gene compartments can drive condensates to reposition preferentially to a single compartment, adopt elongated morphologies, or undergo fission. Together, our model provides a unified framework to explain diverse properties and puzzling observations underlying nuclear condensates. More generally, our model provides a step towards advancing our understanding of how non-equilibrium processes impact regulation and dynamics of multicomponent condensates and genome organization.

## Results

### Model of active nuclear condensates

Many nuclear condensates enrich molecules that catalyze gene expression ^20,25,37^ and are proximate to sites of transcription on the genome. The genome itself is organized into spatially clustered hubs of active and inactive genes, referred to as A and B compartments, arising through structural and sequence-based interactions amongst the polymeric DNA scaffold, nuclear proteins, and RNA^9,12,23^. Actively transcribed genes, in turn, modulate both genomic compartments and properties of nuclear condensates in an RNA-dependent manner^11,25,30,52^. How this complex interplay between chromatin compartmentation, active RNA synthesis, and multiple nuclear condensates dictate organization and function is poorly understood.

To explore how genome activity couples to nuclear condensates, we developed a coarse-grained physics-based model (Figure 1; SI Model) that builds on our previous work^27^. In this model, genomic compartmentation into active hubs is effectively described by a spatially clustered region of gene activity (Figure 1 – green plots). We model a single active compartment of genes (SI Model), described by their total transcriptional activity 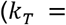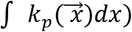 and extent of spatial clustering or compartmentation (*σ*.). Highly expressed genes contribute to higher *k*_*T*_ while stronger spatial clustering of genes reflect a smaller *σ*. Common to these condensates are high local concentrations of nucleic acids and proteins that often catalyze transcription. To capture this, we employ a coarse-grained description in which protein and RNA components are each modeled as an effective pseudospecies (Figure 1, blue and pink species). Attractive interactions between protein and RNA components promote condensation at particular stoichiometries while preferring the soluble (or mixed) phase at asymmetric stoichiometries (Figure – equations for free-energy). This model is motivated by an electrostatic complex-coacervate model of transcriptional condensates that we previously developed^27^ but extends to condensates whose assembly is primarily driven by *heterotypic* interactions, as is the case with many biological condensates^53^. The protein and RNA components diffuse with specified mobility coefficients (Equations, Figure 1). RNAs are actively synthesized at a rate depending on the density of transcriptional proteins as well as genomic activity (Figure 1, SI Model) and degraded at a uniform rate. The evolution of spatiotemporal dynamics of this system, as described by the RNA and protein concentrations (*ϕ*_*R*_(*x*, *t*), *ϕ*_*P*_(*x*, *t*)), is simulated through evolving a continuum phase-field model on a 2D circular grid (Figure 1 – equation for dynamics). Unless specified otherwise, the initial conditions begin with uniform RNA background and a condensate nucleus proximate to the site of genomic activity (SI Simulations). The simulation data is analyzed (SI Analyses) to obtain measurements of condensate size, stability, morphology, and dynamics.

**Figure 1.**
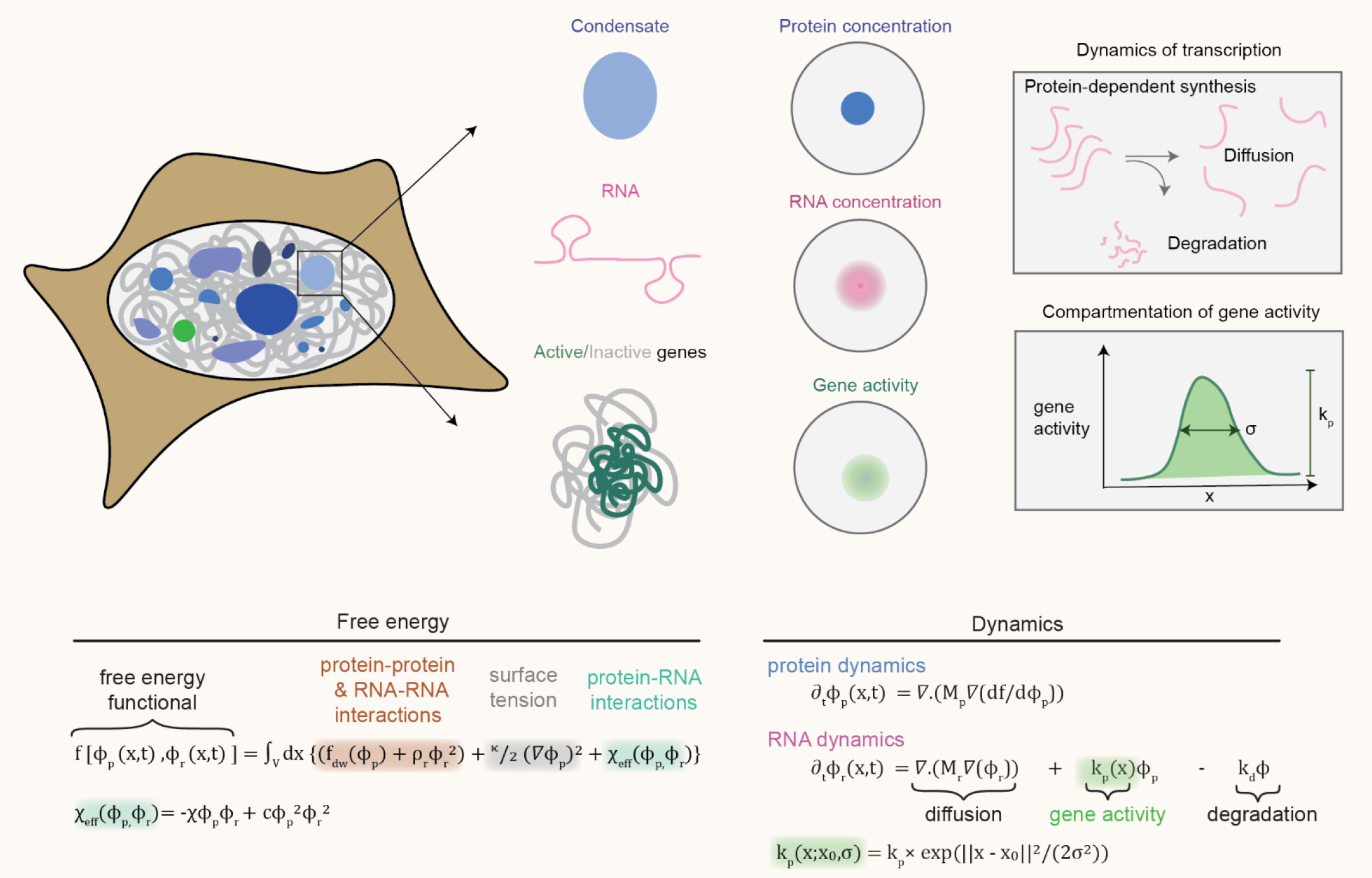
Model of active nuclear condensates. Nuclear condensates are characterized by a dense phase of proteins, which are also often enriched in RNA species. These condensates can overlap with regions of high gene activity. The rate processes of RNA synthesis from these genes, degradation, and diffusion shape the dynamics and organization of nuclear condensates. Below the schematic, the equations used to model the free-energy as well as the coupled dynamics of proteins & RNA are specified.

### Spatial clustering of gene activity dictates condensate size and nucleation

Across a wide range of cell-types and organisms, transcribed genes are spatially clustered in the nucleus ^9,52,54^ (also known as A-type compartments) and often found adjacent to specific nuclear condensates^31,40^. How and whether compartmentation of gene activity influences condensate localization and dynamics in the nucleus is not known. To explore how compartmentation influences condensate properties, we ran simulations contrasting scenarios where genes are clustered (*σ* = 2) or uniformly distributed (Figure 2A) and varying the RNA synthesis rate (*k*_*T*_) while holding other parameters constant. We find that increasing gene activity first promotes, and subsequently dissolves, active nuclear condensates (Figures 2A-B, S1A) – consistent with our previous findings for transcriptional condensates ^27^. For the same total rate of RNA synthesis *k*_*T*_,we find that clustered genes encode for higher local concentrations of RNA compared to a spatially uniform gene density, which in turn, shifts the regime of condensate stability to lower transcription rates (Figure 2A; black line – clustered, grey line - uniform). Spatial clustering of genes is sufficient to recapitulate this phenomenon, irrespective of the coarse-grained representation of cluster shape (Figure S1B). Reducing density or compartmentalization of active genes i.e. increasing *σ*, has a more modest effect on condensate size (Figures 2C-D, Figure S1C). Physically, reducing the degree of compartmentalization i.e. gene clustering (increasing *σ*) leads to lower local concentrations of RNA. Thus, decreasing clustering leads to diffuse RNA concentrations over larger regions leading to larger condensates (Figure 2C). At very low compartmentation (very high *σ*), condensates stop growing and become smaller. This happens because when RNA concentrations are diffuse, the low concentrations are insufficient to recruit protein apparatus (Figure 2C). Together, our model shows that both activity and extent of spatial clustering of genomically active regions modulates size and stability of nuclear condensates. Importantly, genes that are spatially clustered can promote condensate stability even at low levels of transcription. This may explain why nuclear condensates often form around clustered genomic regions with wide range of transcriptional activities, such as super-enhancers (transcriptional condensates), rDNA repeats (nucleoli), and histone-gene repeats (HLBs). Consistent with emerging experimental studies, our model provides a mechanistic framework that implicates RNA and genome structure as key players in regulating nuclear condensate organization *in cis* ^24,25,55,56^.

**Figure 2.**
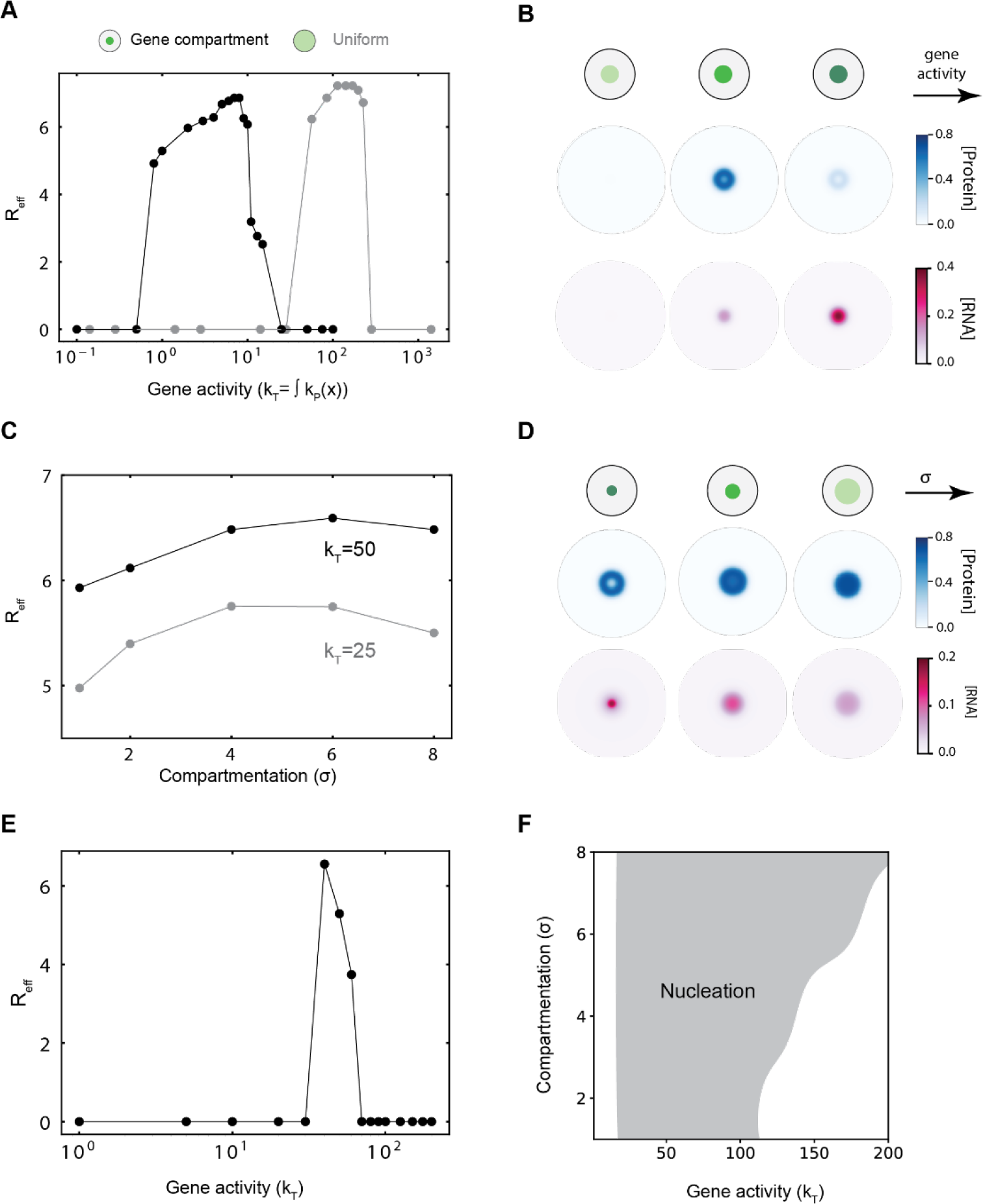
Spatial clustering of gene activity dictates condensate size and nucleation. **A.**Condensate radius (*R*_*eff*_) plotted against gene activity (*k*_*T*_) for two different cases: (i) the gene activity is localized to a region in space with a compartmentation value of *σ* = 2 (black curve) (ii) the gene activity is constant in space (grey curve). The gene activity here is defined as the is the cumulative rate constant over space (*k*_*T*_ = ∫ *k*_*P*_(*x*).2*πx*. *dx*). For these simulations, a dense phase of protein was nucleated at the center of the simulation domain in a background of dilute protein. **B.**Steady state concentration profiles of protein (blue) and RNA (pink) as we increase gene activity *k*_*T*_ from left to right. **C.**Condensate radius (*R*_*eff*_) plotted against the extent of compartmentation (*σ*) for two different values of *k*_*T*_. A small value of *σ* corresponds to highly localized gene activity while *σ* → ∞ corresponds to uniform gene activity **D.**Steady state concentration profiles of protein (blue) and RNA (pink) as we increase *σ* (decrease localization) from left to right **E.**Radius (*R*_*eff*_) of condensate nucleated by activity alone as a function of gene activity (*k*_*T*_). For these simulations, the initial condition is a uniform dilute protein concentration everywhere. No dense phase of protein was nucleated at the center of the simulation domain. **F.**Phase diagram of regions where a condensate get nucleated due to gene activity, upon varying its magnitude (*k*_*T*_) and compartmentation (*σ*).

Nuclear condensates are subject to dynamic control across the cell cycle and often form around specific genomic loci ^6,37,57^. To explore whether genomic compartments can drive assembly of nuclear condensates *de novo*, we ran simulations where no condensates are initially present in the simulations (Figure 2E; SI Simulations). Upon increasing the total activity of the genomic compartment, the model predicts *de novo* condensate nucleation at the site of active transcription (Figure 2E). Nucleation requires spatially localized concentrations of RNA and does not occur e upon removal of genomic compartmentation (Figure S1D; left panel). Further, kinetics of nucleation are faster at moderate rates of transcription (Figure S1D; right panel). By simulating a range of gene activities and extent of compartmentation, our model predicts that moderately clustered regions of gene activity are sufficient to drive local nucleation of condensates (Figure 2F). Together, this model provides a plausible basis to explain diverse observations of transcription-dependent nucleation of condensates, with prominent examples including paraspeckles^33,34,38^, nuclear-stress bodies^33,34^, and specific nucleolar layers^37^.

### Active nuclear condensates exhibit unusual steady-state morphologies

Condensates exhibit diverse morphologies in cells^5,38,39^ unlike the symmetric spherical morphology expected from models of simple Newtonian two-phase fluids. While this discrepancy has been partly ascribed to the viscoelastic nature of many condensates^58^, how non-equilibrium processes affect shape is not well described. This is particularly relevant in the nucleus where complex morphologies are often lost upon inhibition of active transcription^39,42,59^.

To explore how gene expression modulates condensate morphology, we first expanded upon our observation of vacuole formation at high transcription rates (Figures 2B,2D). Vacuoles are regions of low protein but high RNA that form within an otherwise protein-rich condensate (Figure 3A bottom panel). Vacuoles form due to the RNA/prote in rich demixed phase becoming locally unstable due to high RNA concentrations that arise from clustering of highly active genes. These vacuoles are lost upon lowering activity (Figure 3A, top panel) and have dual interfaces (det(J)<0 in Figure 3A; Methods) between an inner RNA-rich protein-poor and outer RNA/protein-poor phases - representing a non-equilibrium core-shell morphology. By simulating a range of activities and compartmentation (*k*_*T*_, *σ*), we find that high activity and moderate/strong compartmentation (low *σ*) is required for vacuole formation (Figures 3B, S2A). Physically, at low activity or weak clustering (low *k*_*T*_, high *σ*), RNA concentrations are not locally high enough to create vacuoles and very high activity or compartmentation completely dissolves condensates – thus, vacuoles form at intermediate activity and compartmentation under simulation conditions. Nucleoli, condensates built around highly clustered rDNA gene repeats and transcribed at high levels have been previously documented to contain vacuoles^13,42^ and exhibit a multilayered condensate structure, with transcription concentrated within the central regions^39^. Layered morphologies are lost or modified upon inhibition of transcription^39^, suggesting an important role for activity. Our model’s prediction of vacuoles is broadly consistent with these observations. While prevailing models emphasize the importance of biomolecular interactions^39,60,61^ and dynamic processes beside transcription^60,62^, our study implicates transcriptional activity as an additional axis regulating nucleolar organization.

**Figure 3.**
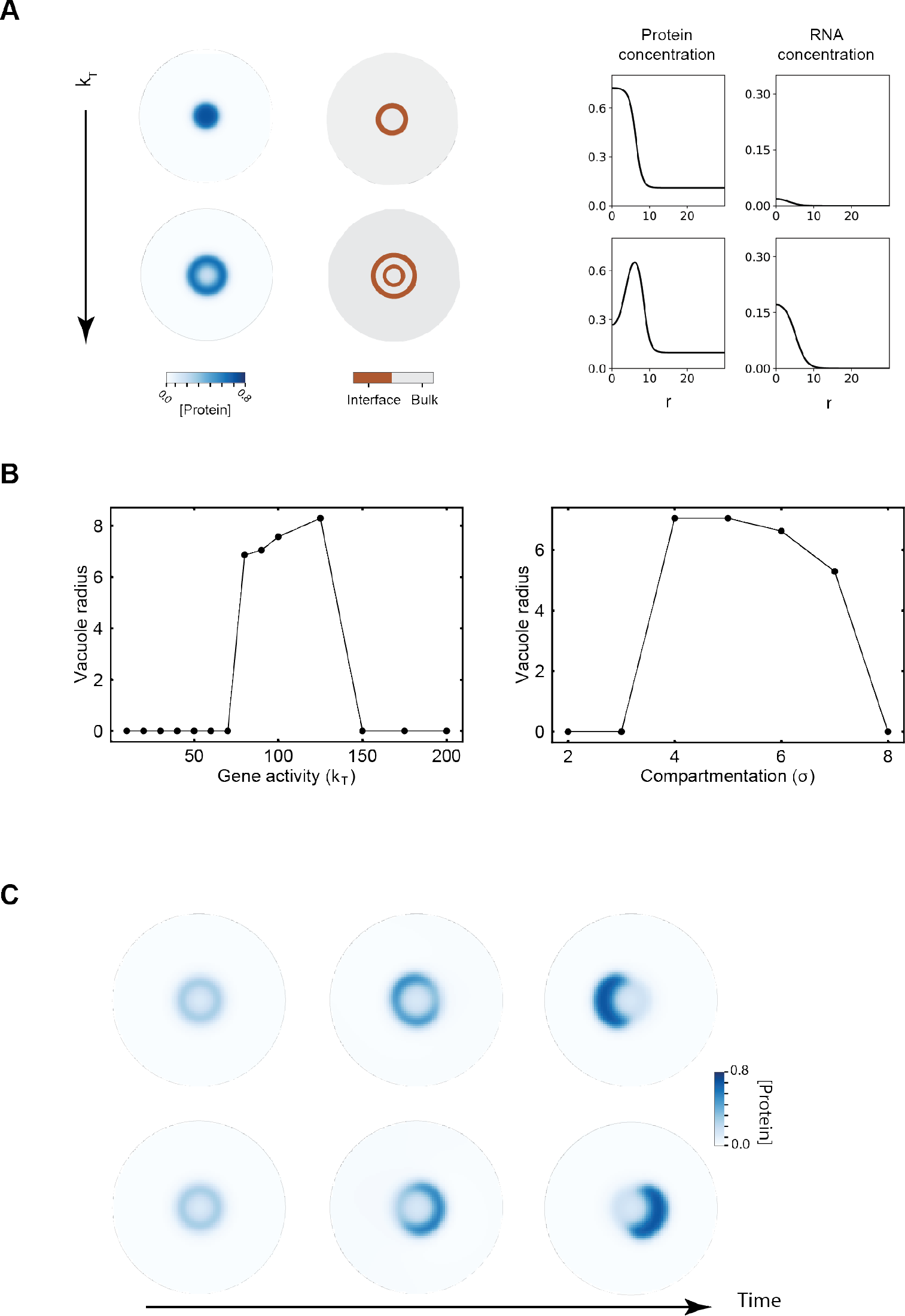
Active nuclear condensates exhibit unusual steady-state morphologies. **A.** In the left panel, the blue plots correspond to the protein concentration profiles and the brown plots represent interfaces between two phases for two different values *k*_*T*_. The right panel plots concentration profiles of the protein and the RNA as a function of the radial coordinate *r*. In the top panel with *k*_*T*_ = 1.0 (*σ* = 3), we have a single dense phase of protein separated from the dilute phase by a single interface. In the bottom panel, we have a ring-like dense phase of protein sandwiched between a dilute phase inside and outside, each separated by respective interface. We call the dilute phase inside the ring as the vacuole. Increasing the gene activity to *k*_*T*_ = 20.0 (*σ* = 3) can lead to a change in condensate morphology from a single droplet (top panel) to a vacuole (bottom panel). **B.** Radius of the vacuole (the dilute phase of protein within the ring) plotted against different magnitudes of gene activity *k*_*T*_ (with *σ* = 4 fixed) and extents of compartmentation *σ* (with *k*_*T*_ = 90.0 fixed) **C.**Protein concentration profiles (blue) as time progresses for two different initial conditions where *k*_*T*_ is large. Both simulations are done for *k*_*T*_ = 50.0 and *σ* = 4. We observe that a symmetric vacuole is initially formed. This symmetry is broken as time progresses and we end up with an aspherical droplet.

At high transcription rates, we find that vacuoles spontaneously break symmetry to form aspherical condensates that partially overlap the site of active transcription (Figure 3C). Physically, larger activities increase vacuole size, which in turn, increases surface costs due to dual interfaces. This eventually leads condensates to adopt aspherical morphologies with lower surface areas to minimize surface tension costs, even transiently forming 2 aspherical condensates under certain parameter conditions (Figure S2C). Condensates prefer to partially overlap the interface of the active site rather than break into smaller spherical droplets. Physically, this is because spatially decreasing activity and RNA concentration leads to favorable interactions at the interface of the gene that stabilize condensation but surface tension costs limit stability of spherically symmetric vacuoles – leading to adoption of aspherical morphologies. Consistent with this logic, increasing the surface tension at fixed activity in our simulations leads to vacuoles breaking symmetry before eventually dissolving due to high interface costs and vice-versa (Figure S2D). Condensates break symmetry at different positions that are uniformly distributed around the gene across multiple trajectories with slightly different initial conditions, indicating no preferred direction (Figure S2B). Since vacuole formation and aspherical morphologies depend on high local RNA concentrations, we reasoned that increasing the mobility of RNA while holding parameters constant should lower local RNA concentrations. Note that such a perturbation is purely dynamic in nature and does not affect equilibrium properties. We find that lower effective concentrations (or higher *M*_*rna*_) leads to a transition from dissolved condensates to aspherical morphologies and eventually vacuoles (Figure S2E), highlighting the importance of dynamics and transport in driving these non-equilibrium morphologies. Overall, our results suggest that strong compartmentation and high transcriptional activity gives rise to non-equilibrium morphologies like vacuoles and aspherical droplets. This may underlie why nuclear speckles, which are condensates that experience a high RNA flux, often adopt *granular* morphologies which subsequently become spherical upon inhibition of gene expression ^59,63^ (or loss in RNA flux). Aspherical condensates localized to the edge of active sites are observed in our models, mimicking the interfacial localization of actively transcribed genic regions around nuclear speckles^31,64^ - although additional factors likely play important roles to drive *in vivo* organization, including interactions between transcriptional and splicing proteins as well as post-translational modifications^64,65^.

### Distant gene activity induces flow of nuclear condensates

Condensates are present in crowded and heterogeneous environments in the cell and typically move randomly, often sub-diffusively^46–48,66^, limited by the chromatin or cytoskeletal network. However, many condensates also exhibit bursts of super-diffusive or directed motion^47,48^, particularly upon signaling or stress^49^. A mechanistic basis for this directed motion remains lacking.

To explore long-range condensate motion, we developed simulations where the active site and gene condensate are initially separated by a distance *r*. Note that our model neglects thermal diffusion of condensates, which are often slow^66^ in cells i.e. minutes-long time-scales, and any motion observed is purely directional within these short time scales. We found that condensates that are initially located far away from the active site flow towards it and upregulate transcription (Figure 4A). Directed motion occurs only when the condensate is closer than a distance *r*^*^ from the active site, and if further away, the condensate does not move (Figure 4B). Since there is low amounts of transcription (mimicking basal gene expression) at the active site in the absence of a condensate, we wondered if that led to a gradient in RNA which in turn drove condensate motion. To explore this, we computed an estimate of the length-scale of gradient i.e. spatial extent of RNA concentration, from the ratio of reactive and diffusive length-scales 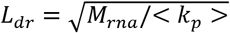. This length scale measures how far RNA molecules spread, with higher mobility (*M*_*rna*_; units of 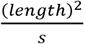) leading to more spread-out distributions (higher *L*_*dr*_) and faster rates of transcription (higher *k*_*p*_; units of 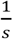) leading to quicker turnover before diffusion, which in turn, leads to a contained spatial distribution of RNA (lower *L*_*dr*_). When this gradient length-scale was greater than the initial separation between the gene cluster and condensate (*L*_*dr*_/*r* > 1), our simulations showed that the condensate moved towards the active site and when the gradient was shorter, there was no directed motion (Figure 4B). If gradient-sensing was the mechanism underlying directed motion, we reasoned that condensates that are far away can pick up an RNA gradient if RNA mobility was larger. As predicted, increasing *M*_*rna*_, and thus *L*_*dr*_, led to directed motion beyond a threshold mobility - importantly, this threshold approximately corresponded to a gradient whose length-scale matched the initial separation (Figure 4C). For a fixed transcription rate, increasing mobility increases *length* of gradient by distributing a similar amount of content over larger scales, which means the *strength* (or magnitude) of gradient, and thus the velocity of motion would be smaller. Consistent with these expectations, we found that stronger but shorter gradients (low *M*_*rna*_) led to faster flow than weaker but longer gradients (Figure S3A). Condensates with higher surface tension moved at similar speeds as those with lower surface tension but underwent lesser distortion, as measured by the change in eccentricity over time (Figure S3B). Overall, our model predicts that if a site of active transcription generates a gradient strong enough to be “sensed” by the condensate (*L*_*dr*_ /*r* >≈ 1), this leads to directed flow. Based on typical rates of RNA diffusion and transcription in cells (SI Gradient Calculation), which can intrinsically vary significantly^67–69^, we estimate that gradients in RNA fluxes potentially span a range of scales from about 0.1 − 1*μm*. These length-scales are comparable to condensate sizes^20^, and thus, experimentally discernible if driving short bursts of directed motion. While our model neglects thermal diffusion, active nuclear condensates that are otherwise diffusing randomly can exhibit directional motion when a nearby gene compartment becomes active. This provides a mechanistic framework to explain why different nuclear bodies exhibit *super-diffusive* or partly ballistic motion in their cellular trajectories^46–48^ and may provide a mechanism for how nuclear bodies quickly and dynamically allocate machinery upon signaling or stress.

**Figure 4.**
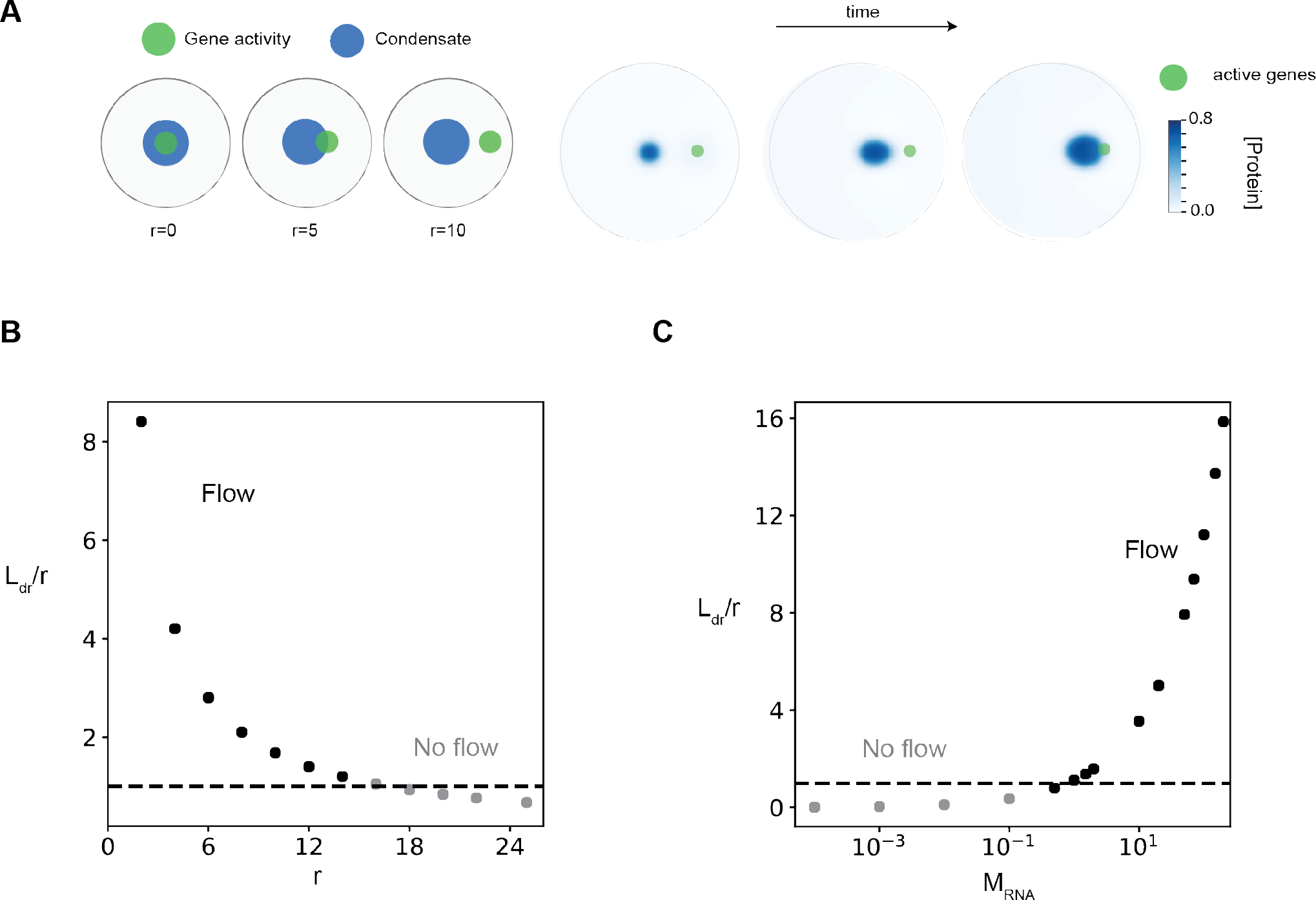
Distant gene activity induces flow of nuclear condensates. **A.** The panel is a cartoon illustrating the initial condition where the center of the condensate (blue) is located at different distances (*r*) from the center of the site of gene activity (green). The right panel illustrates a condensate (blue) growing and moving towards a site of active genes (green) as time progresses, for the case where *r* = 10. For these simulations, the parameters used are *k*_*T*_ = 10.0, *σ* = 4, and *M*_*RNA*_ = 1.0 **B-C.** The ratio of RNA diffusion length scale (*L*_*dr*_) to the initial distance between condensate (*r*) and site of gene activity for different values of *r* and the RNA mobility *M*_*RNA*_. The RNA diffusion length scale is calculated as *L*_*dr*_ = (*M*_*RNA*_/< k_p_ >)^1/2^, where < *k*_*p*_ > is the average activity. Decreasing the distance *r* for a fixed value of *L*_*dr*_ leads to change in phenotype of condensate dynamics from a case where there is no flow to one where the condensate flows towards the site of gene activity. Similarly, increasing the RNA mobility (*M*_*RNA*_) leads to a change from No flow → Flow. In both cases, the ratio of the length scales *L*_*dr*_ /*r* crossing the value of 1.0 predicts the transition from No flow → Flow.

### Gene activity and position dictate emergent condensate morphology and dynamics

Given the diverse morphology and dynamics we observed, we sought to derive a *phase* space of possible outcomes with varying compartment activity and relative initial position of condensate (Figure 5). We found that condensates were confined to the gene compartment when initially close by (Figure 5, top panel) irrespective of transcriptional activity. Condensates responded to the presence of a distant active gene compartment by dissolving and subsequently nucleating at the gene compartment, but only when the transcriptional activity was high enough (Figure 5, top panel). At intermediate distances, condensate dynamics was governed by directed flow (Figure 5, top panel). These data suggest that dynamics of condensates vary, undergoing directed motion when close by but not adjacent to a gene compartment, and leading to activity-dependent long-range dissolution and nucleation when present far away. Physically, when RNA concentrations are diffuse, condensates that partially overlap flow up these gradients. However, when condensates are sufficiently far away from a highly active gene cluster, the cluster promotes local nucleation of a condensate. Since overall protein concentrations are fixed in our simulations, motivated by their slow turn-over in cells, this led to a concordant dissolution of the distant condensate. By contrast, the steady-state morphologies of the condensate were nearly independent of initial positions. With increasing gene activity, spherical droplets developed vacuoles, eventually leading to symmetry-breaking (see Figure 3) and adoption of aspherical morphologies (Figure 5, bottom panel). At very low activities, when the condensates are far from gene clusters, they do not colocalize with clusters as they do not sense the gradient. The primary exception to this lack of dependence on initial positions occurs when condensates are nucleated around high activity genic compartments, leading to their dissolution before repositioning or adoption of different morphologies. Overall, our model partitions condensate dynamics and morphology into two axes - distance from active gene clusters primarily modulates dynamics while transcriptional activity mostly dictates steady-state morphology. This model provides a framework to potentially explain why condensates around regions of high activity, such as nucleoli or nuclear speckles, often exhibit non-equilibrium and aspherical morphologies^30,39,49,59^, provides mechanisms by which nuclear bodies exhibit directed or super-diffusive motion^46–48^, and shows that compartmentalized gene activity may be sufficient to locally nucleate a condensate^33,34,70^.

**Figure 5.**
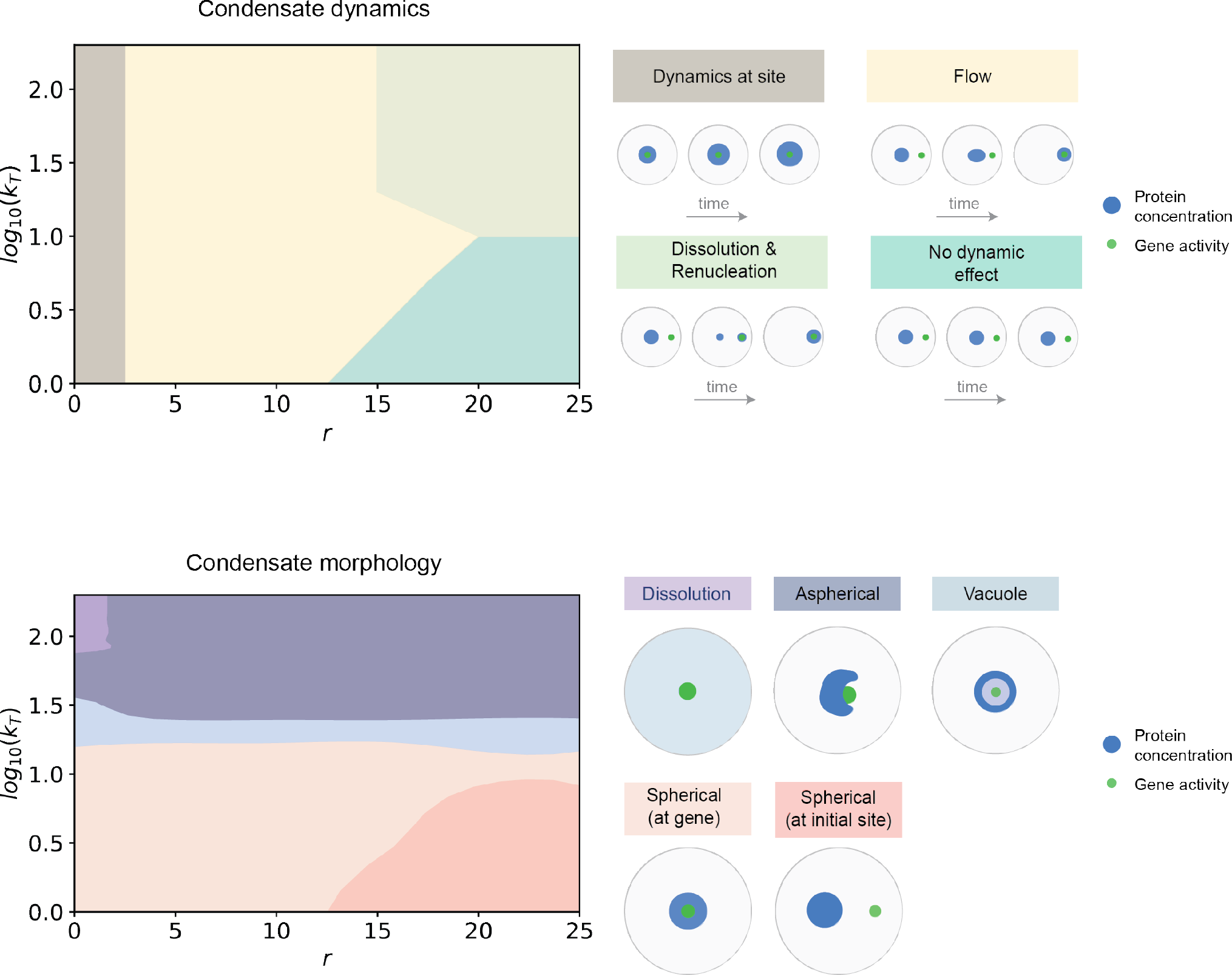
Gene activity and position dictate emergent condensate morphology and dynamics. The top panel summarizes a diagram of the different qualitative dynamics - dynamics at site, flow, dissolution & renucleation, no effect - that are possible when varying distance *r* and total gene activity *k*_*T*_. The bottom panel summarizes a diagram of different qualitative morphologies - spherical, aspherical, vacuole, dissolution - that are possible upon varying the distance *r* and the magnitude of gene activity *k*_*T*_. All simulations were done with *σ* = 4.

### Multiple genic compartments compete for or share nuclear condensates

The nucleus of a cell is highly heterogeneous, filled with many different compartments with varying gene activity and multiple condensates^31,32,52^ that often interact across large distances. While our primary focus has been to dissect the interplay between nuclear condensates and a single active gene compartment, we next sought to explore how multiple active compartments modulate condensates. To this end, we ran simulations with 2 active compartments, whose properties (activity, extent of clustering or gene density, and position) we changed while holding other parameters constant. Note that we held the total protein levels constant and limiting for all cases. We first considered 2 nearby compartments - one active (gene cluster B – right compartment in Figure 6A) and one whose activity we varied (gene cluster A – left compartment in Figure 6A). In the absence of or at low levels of activity in cluster A, nuclear condensates colocalize with gene cluster B at steady-state (Figure 6A). As the activity of cluster A is increased, the condensate repositions to overlap both sites (Figure 6A, ratios=1, silver line), and when gene cluster A’s activity is higher, the condensate adopts an aspherical morphology centered on cluster B but interfacially localizing around cluster A, similar to Figure 3C. In the latter cases, as activity of gene cluster A is very high, condensates don’t completely overlap with the high RNA concentrations. We next explored how changing compartmentation, but not total activity, of gene cluster A influences condensate morphology. When gene compartments are similar in size but apart by a finite distance *r*, condensates adopt a stretched or elongated morphology to overlap both sites. Decreasing compartmentation (increasing *σ*_*A*_) lessens the local RNA concentrations from cluster A, and so the condensate moves to occupy gene cluster B (Figure 6B). Conversely, increasing compartmentation (decreasing *σ*_*A*_) increases local RNA concentrations, moving the condensate to cluster A (Figure 6B). Rather than changing gene features, if we vary the relative position of two genetic clusters, we find that condensates initially elongate to accommodate both sites but eventually splinter to one of two sites (Figure 6C). This splitting occurs in part due to protein levels being limited. When protein levels are higher, condensates split into 2 droplets rather than elongate when surface tension costs are low (Figure S4). Overall, these results suggest that a combination of compartment strength and activity, as well as their relative positions in the nucleus, contribute to organization and morphology of condensates. When both gene compartments are similar and are active, our model finds that condensates reposition and often elongate to overlap both sites, contributing to aspherical morphologies. This may underlie why RNA Pol II condensates, which often span multiple enhancers and genetic loci, exhibit elongated or unfolded shapes in cells^71^, as well as other multi-compartment spanning condensates like speckles^33,59^.

**Figure 6.**
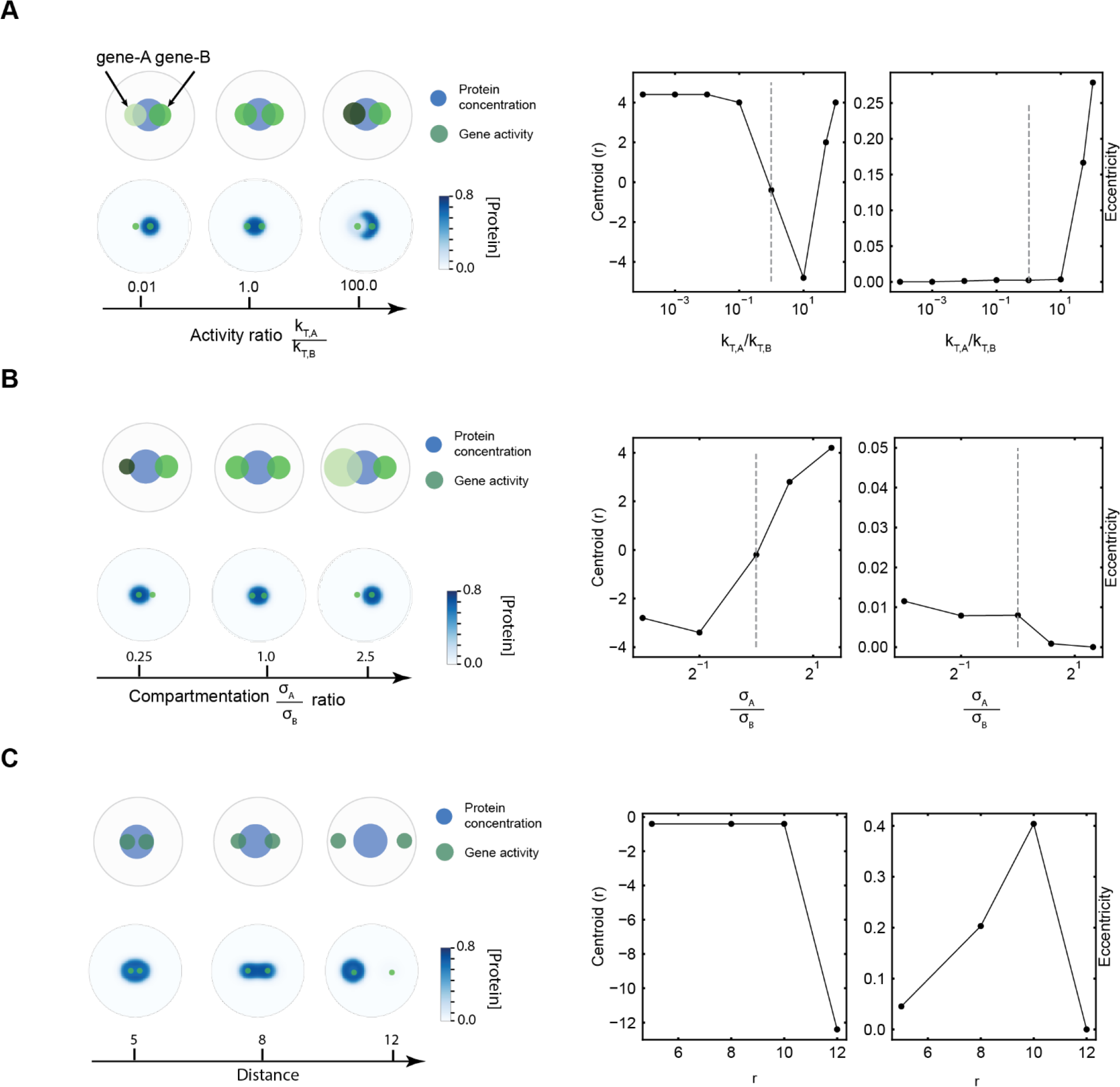
Multiple genic compartments compete for or share nuclear condensates. **A.** The ratio (*k*_*T*,*A*_/*k*_*T*,*B*_) of gene activity between gene-A and gene-B is increased by increasing the magnitude of gene activity *k*_*T*,*A*_ from 10^−4^ to 100 while keeping *k*_*T*,*B*_ = 1.0 fixed. The steady state concentration profile of the protein is shown in blue for different values of the ratio (*k*_*T*,*A*_/*k*_*T*,*B*_). The right panel plots the centroid and eccentricity of the dense phase of protein for different values of the ratio (*k*_*T*,*A*_/*k*_*T*,*B*_). Refer SI for more details on how the centroid and eccentricity are calculated. A value of *σ* = 4 was used for both gene-A and gene-B. **B.** The ratio of compartmentation (*σ*_*A*_/*σ*_*B*_) of gene activity is increased by increasing *σ*_*A*_ from 1.0 to 10.0, while keeping *σ*_*B*_ = 4.0 fixed. The steady state concentration profile of the protein is shown in blue for different values of the ratio (*σ*_*A*_/*σ*_*B*_). The right panel plots the centroid and eccentricity of the dense phase of protein for different values of the ratio (*σ*_*A*_/*σ*_*B*_). A value of *k*_*T*_ = 10.0 was used for both gene-A and gene-B. **C.** The steady state concentration profile of the protein is shown in blue for different values of distance (*r*) between the centers of gene-A and gene-B. The right panel plots the centroid and eccentricity of the dense phase of protein for different values of *r*. A value of *k*_*T*_ = 10.0 and *σ* = 4 was used for all simulations.

## Discussion

Condensates are biomolecular assemblies that compartmentalize and organize active cellular processes. In the nucleus, condensates interact with the genome and often regulate active gene expression through concentrating specific molecules or pathways1,20. The genome, in turn, is highly organized in the cell, most prominently into territories of individual chromosomes at large length-scales, and sub-divided into many smaller compartments that are broadly enriched in either active or inactive genes^9,11,12^. It is quickly emerging that the interplay between genome organization and nuclear condensates occurs in an RNA/transcription-dependent manner^5,25,27^ and is important for function. Dissecting this complex coupling has remained challenging, in part due to the lack of model frameworks. In this study, we develop a physics-based model to investigate how gene compartments (or hubs) interact with nuclear condensates that form by phase separation, and the role of active transcription (RNA synthesis) in modulating emergent properties of condensate dynamics, morphology, and organization.

First, we develop a coarse-grained model of the underlying process with 3 main features (1) Description of nuclear condensate phase behavior in terms of RNA/protein interactions and concentrations (2) Spatial model of gene compartments and (3) Dynamical coupling between condensates and gene compartments through direct activation of transcription when proximate (Figure 1). Through simulating a range of conditions where features of gene compartment (activity, extent of compartmentation) are varied, we identify that even low levels of activity, when clustered spatially, stabilize nuclear condensates (Figure 2). Across a broad range of activities and compartment sizes, we find that gene clusters directly and specifically nucleate condensate formation (Figure 2). Overall, these results show that activity and compartment strength directly influence condensate size and dynamics. These results provide a mechanistic basis to interpret why distinct nuclear condensates assemble specifically around clustered genomic elements like super-enhancers^21,72^, histone-gene repeats^26,36,57^, or rDNA repeats^39^ and explain how multiple condensates may nucleate in a transcription-dependent manner^34,35,70^. While our model is coarse-grained into describing effective RNA and protein species, an exciting area of research will be to explore how the diverse network of interactions between specific proteins, regulatory DNA elements, and RNA ^5,21,61,73^ shape gene regulation, RNA processing, and nuclear organization for distinct condensates. Although we focused on condensates that activate transcription, similarly motivated models will contribute to better understanding of the emerging nexus between inactive heterochromatin and RNA-dependent nuclear condensates^35,74–76^ as well as interactions between different types of condensates.

We find that the presence of compartmentalized gene activity, albeit at high levels, gives rise to non-equilibrium steady-state morphologies such as vacuoles and aspherical condensates (Figure 3). These arise due to high RNA concentrations that locally disfavor the demixed phase and are not observed if activity is lowered. Aspherical or layered morphologies have been previously described for multiple nuclear condensates that abut highly transcribed loci and are sensitive to inhibition of transcription^29,39,43^. Our model provides a mechanistic basis to explain how *active* RNA fluxes can drive emergent condensate morphologies and likely contribute to morphology *in vivo*. Through simulations, we next find that condensates can “sense” and exhibit directed motion towards distant compartments (Figure 4), limited by transport constraints that we estimate through a simple physical calculation of the RNA gradient. This highlights a potential mechanism by which condensates dynamically sense and allocate machinery to genes upon signaling or stress and betters our understanding of diverse yet mechanistically unexplained observations of super-diffusive and long-range motion in nuclear bodies^46–49^. We classify both dynamic and steady-state features of active nuclear condensates, with activity being the primary determinant of steady-state morphology and initial position driving dynamics (Figure 5). While our model focuses on condensate properties at smaller length and time-scales, an important future area of research will be to dissect how condensates directly restructure chromatin structure over longer-time scales and contribute to large-scale nuclear organization^31,66^,including through introduction of capillary forces that arise from condensates wetting different biological substrates^77^.

Finally, we find that condensates can exhibit activity-dependent repositioning, elongated morphologies, and fission when a second active compartment is introduced in the vicinity of the first (Figure 6). These predictions provide a plausible explanation for the organization of Pol II clusters and nuclear speckles, condensates that typically span multiple active sites, into elongated unfolded morphologies^59,71^ and observations of condensate fission^47^. Motivated by observations and puzzles in nuclear condensate phenomenology, our model provides a unified framework to interrogate how the interplay between genomic compartments, active transcription, and phase separation influence condensate stability, dynamics, and morphology. Emerging techniques that directly engineer and visualize condensates at specific genomic loci^78–80^ provide exciting avenues to test and refine existing models. In turn, new theories and models will likely be required to investigate how non-equilibrium processes influence multicomponent multiphase fluids^81–83^. How active processes in general, including chemical fluxes and ATP-dependent molecular motors, couple to genome organization and condensate properties in normal and pathological states represents an important frontier of future research.

## Materials

Phase-field simulations and subsequent data analyses were performed using custom code written in python and salient aspects of the model are briefly described in Figure 1. For a detailed description of the model, simulation, and analyses, please refer to the accompanying Supplementary Information.

## Supporting information

Unembedded_main_figures

## Acknowledgments

K.S. acknowledges support from the NSF–Simons Center for Mathematical and Statistical Analysis of Biology at Harvard (Award #1764269) and the Harvard Faculty of Arts and Sciences Quantitative Biology Initiative. P.N. and A.K.C acknowledge support from NSF (Award #2044895).

## Author contributions

K.S. and A.K.C conceived the project. H.H.S. and K.S. developed the model. H.H.S and P.N ran simulations and analyzed data. K.S. designed figures with the help of P.N. and H.H.S. K.S. wrote a first draft and all authors contributed to editing and revising the manuscript.

## Code

The code for running simulations is available at https://github.com/npradeep96/RNA_localization_final

## References

1. Banani, S. F., Lee, H. O., Hyman, A. A. & Rosen, M. K. Biomolecular condensates: organizers of cellular biochemistry. Nat. Rev. Mol. Cell Biol. 18, 285–298 (2017).

2. Lyon, A. S., Peeples, W. B. & Rosen, M. K. A framework for understanding the functions of biomolecular condensates across scales. Nat. Rev. Mol. Cell Biol. 1–21 (2020) doi:10.1038/s41580-020-00303-z.

3. Shin, Y. & Brangwynne, C. P. Liquid phase condensation in cell physiology and disease. Science 357, eaaf4382 (2017).

4. Choi, J.-M., Holehouse, A. S. & Pappu, R. V. Physical Principles Underlying the Complex Biology of Intracellular Phase Transitions. Annu. Rev. Biophys. 49, (2020).

5. Roden, C. & Gladfelter, A. S. RNA contributions to the form and function of biomolecular condensates. Nat. Rev. Mol. Cell Biol. 1–13 (2020) doi:10.1038/s41580-020-0264-6.

6. Rai, A. K., Chen, J.-X., Selbach, M. & Pelkmans, L. Kinase-controlled phase transition of membraneless organelles in mitosis. Nature 1 (2018) doi:10.1038/s41586-018-0279-8.

7. Berry, J., Brangwynne, C. P. & Haataja, M. Physical principles of intracellular organization via active and passive phase transitions. Rep. Prog. Phys. (2018).

8. Weber, C. A., Zwicker, D., Jülicher, F. & Lee, C. F. Physics of active emulsions. Rep Prog Phys 82, 64601 (2019).

9. Mirny, L. & Dekker, J. Mechanisms of Chromosome Folding and Nuclear Organization: Their Interplay and Open Questions. Cold Spring Harb. Perspect. Biol. a040147 (2021) doi:10.1101/cshperspect.a040147.

10. Lieberman-Aiden, E. et al. Comprehensive Mapping of Long-Range Interactions Reveals Folding Principles of the Human Genome. Science 326, 289–293 (2009).

11. Misteli, T. The Self-Organizing Genome: Principles of Genome Architecture and Function. Cell 0, (2020).

12. Bonev, B. & Cavalli, G. Organization and function of the 3D genome. Nat. Rev. Genet. 17, 661–678 (2016).

13. Balbiani, E. G. Sur la structure du noyau des cellules salivaires chez les larves de Chironomus. Zool Anz 4, 662–667 (1881).

14. Wagner, R. Einige bemerkungen und fragen über das keimbläschen (vesicular germinativa). Müller’s Arch. Anat Physiol Wiss. Med 268, 373–377 (1835).

15. Wilson, E. B. The structure of protoplasm.*. Science 10, 33–45 (1899).

16. Montgomery, T. H. Comparative cytological studies with especial regard to the morphology of the nucleolus. vol. 5 (Ginn, 1900).

17. Cajal, S. R. & others. Un sencillo metodo de coloracion seletiva del reticulo protoplasmatico y sus efectos en los diversos organos nerviosos de vertebrados e invertebrados. Trab Lab Invest BiolMadrid 2, 129–221 (1903).

18. Gall, J. G. Cajal Bodies: The First 100 Years. Annu. Rev. Cell Dev. Biol. 16, 273–300 (2000).

19. Bond, C. S. & Fox, A. H. Paraspeckles: nuclear bodies built on long noncoding RNA. J. Cell Biol. 186, 637–644 (2009).

20. Sabari, B. R., Dall’Agnese, A. & Young, R. A. Biomolecular Condensates in the Nucleus. Trends Biochem. Sci. 0, (2020).

21. Hnisz, D., Shrinivas, K., Young, R. A., Chakraborty, A. K. & Sharp, P. A. A Phase Separation Model for Transcriptional Control. Cell 169, 13–23 (2017).

22. Sharp, P. A., Chakraborty, A. K., Henninger, J. E. & Young, R. A. RNA in formation and regulation of transcriptional condensates. RNA rna.078997.121 (2021) doi:10.1261/rna.078997.121.

23. Smith, K. P., Hall, L. L. & Lawrence, J. B. Nuclear hubs built on RNAs and clustered organization of the genome. Curr. Opin. Cell Biol. 64, 67–76 (2020).

24. Quinodoz, S. A. & Guttman, M. Essential Roles for RNA in Shaping Nuclear Organization. Cold Spring Harb. Perspect. Biol. a039719 (2021) doi:10.1101/cshperspect.a039719.

25. Bhat, P., Honson, D. & Guttman, M. Nuclear compartmentalization as a mechanism of quantitative control of gene expression. Nat. Rev. Mol. Cell Biol. 1–18 (2021) doi:10.1038/s41580-021-00387-1.

26. Nizami, Z., Deryusheva, S. & Gall, J. G. The Cajal Body and Histone Locus Body. Cold Spring Harb. Perspect. Biol. 2, a000653 (2010).

27. Henninger, J. E. et al. RNA-Mediated Feedback Control of Transcriptional Condensates. Cell 184, 207–225.e24 (2021).

28. Sabari, B. R. Biomolecular Condensates and Gene Activation in Development and Disease. Dev. Cell 55, 84–96 (2020).

29. Chen, Y. & Belmont, A. S. Genome organization around nuclear speckles. Curr. Opin. Genet. Dev. 55, 91–99 (2019).

30. Spector, D. L. & Lamond, A. I. Nuclear speckles. Cold Spring Harb. Perspect. Biol. 3, 1–12 (2011).

31. Takei, Y. et al. Integrated spatial genomics reveals global architecture of single nuclei. Nature 1–7 (2021) doi:10.1038/s41586-020-03126-2.

32. Su, J.-H., Zheng, P., Kinrot, S. S., Bintu, B. & Zhuang, X. Genome-Scale Imaging of the 3D Organization and Transcriptional Activity of Chromatin. Cell (2020) doi:10.1016/j.cell.2020.07.032.

33. Mao, Y. S., Zhang, B. & Spector, D. L. Biogenesis and function of nuclear bodies. Trends Genet. 27, 295–306 (2011).

34. Shevtsov, S. P. & Dundr, M. Nucleation of nuclear bodies by RNA. Nat. Cell Biol. 13, 167–173 (2011).

35. Valsecchi, C. I. K. et al. RNA nucleation by MSL2 induces selective X chromosome compartmentalization. Nature 1–6 (2020) doi:10.1038/s41586-020-2935-z.

36. White, A. E. et al. Drosophila histone locus bodies form by hierarchical recruitment of components. J. Cell Biol. 193, 677–694 (2011).

37. Arias Escayola, D. & Neugebauer, K. M. Dynamics and Function of Nuclear Bodies during Embryogenesis. Biochemistry 57, 2462–2469 (2018).

38. Fox, A. H., Nakagawa, S., Hirose, T. & Bond, C. S. Paraspeckles: Where Long Noncoding RNA Meets Phase Separation. Trends Biochem. Sci. 43, 124–135 (2018).

39. Lafontaine, D. L. J., Riback, J. A., Bascetin, R. & Brangwynne, C. P. The nucleolus as a multiphase liquid condensate. Nat. Rev. Mol. Cell Biol. 22, 165–182 (2021).

40. Quinodoz, S. A. et al. RNA promotes the formation of spatial compartments in the nucleus. Cell 0, (2021).

41. Banerjee, P. R., Milin, A. N., Moosa, M. M., Onuchic, P. L. & Deniz, A. A. Reentrant Phase Transition Drives Dynamic Substructure Formation in Ribonucleoprotein Droplets. Angew. Chem. Int. Ed. 56, 11354–11359 (2017).

42. Johnson, J. M. A study of nucleolar vacuoles in cultured tobacco cells using radioautography, actinomycin D, and electron microscopy. J. Cell Biol. 43, 197–206 (1969).

43. Fei, J. et al. Quantitative analysis of multilayer organization of proteins and RNA in nuclear speckles at super resolution. J. Cell Sci. 130, 4180–4192 (2017).

44. Yasuhara, T. et al. Condensates induced by transcription inhibition localize active chromatin to nucleoli. Mol. Cell 0, (2022).

45. West, J. A. et al. Structural, super-resolution microscopy analysis of paraspeckle nuclear body organization. J. Cell Biol. 214, 817–830 (2016).

46. Görisch, S. M. et al. Nuclear body movement is determined by chromatin accessibility and dynamics. Proc. Natl. Acad. Sci. 101, 13221–13226 (2004).

47. Platani, M., Goldberg, I., Swedlow, J. R. & Lamond, A. I. In Vivo Analysis of Cajal Body Movement, Separation, and Joining in Live Human Cells. J. Cell Biol. 151, 1561–1574 (2000).

48. Platani, M., Goldberg, I., Lamond, A. I. & Swedlow, J. R. Cajal Body dynamics and association with chromatin are ATP-dependent. Nat. Cell Biol. 4, 502–508 (2002).

49. Kim, J., Han, K. Y., Khanna, N., Ha, T. & Belmont, A. S. Nuclear speckle fusion via long-range directional motion regulates speckle morphology after transcriptional inhibition. J. Cell Sci. 132, jcs226563 (2019).

50. Ahn, J. H. et al. Phase separation drives aberrant chromatin looping and cancer development. Nature 1–5 (2021) doi:10.1038/s41586-021-03662-5.

51. Yu, H. et al. HSP70 chaperones RNA-free TDP-43 into anisotropic intranuclear liquid spherical shells. Science 371, eabb4309 (2021).

52. Feric, M. & Misteli, T. Phase Separation in Genome Organization across Evolution. Trends Cell Biol. (2021) doi:10.1016/j.tcb.2021.03.001.

53. Riback, J. A. et al. Composition-dependent thermodynamics of intracellular phase separation. Nature 581, 209–214 (2020).

54. Rowley, M. J. et al. Evolutionarily Conserved Principles Predict 3D Chromatin Organization. Mol. Cell 67, 837–852.e7 (2017).

55. Calandrelli, R. et al. Three-dimensional organization of chromatin associated RNAs and their role in chromatin architecture in human cells. bioRxiv 2021.06.10.447969 (2021) doi:10.1101/2021.06.10.447969.

56. Creamer, K. M., Kolpa, H. J. & Lawrence, J. B. Nascent RNA scaffolds contribute to chromosome territory architecture and counter chromatin compaction. Mol. Cell 81, 3509–3525.e5 (2021).

57. Hur, W. et al. CDK-Regulated Phase Separation Seeded by Histone Genes Ensures Precise Growth and Function of Histone Locus Bodies. Dev. Cell 54, 379–394.e6 (2020).

58. Mittag, T. & Pappu, R. V. A conceptual framework for understanding phase separation and addressing open questions and challenges. Mol. Cell 82, 2201–2214 (2022).

59. Lamond, A. I. & Spector, D. L. Nuclear speckles: a model for nuclear organelles. Nat. Rev. Mol. Cell Biol. 4, 605–612 (2003).

60. Riback, J. A. et al. Viscoelastic RNA entanglement and advective flow underlie nucleolar form and function. 2021.12.31.474660 https://www.biorxiv.org/content/10.1101/2021.12.31.474660v1 (2022) doi:10.1101/2021.12.31.474660.

61. Feric, M. et al. Coexisting Liquid Phases Underlie Nucleolar Subcompartments. Cell 165, 1686–1697 (2016).

62. Wu, M. et al. lncRNA SLERT controls phase separation of FC/DFCs to facilitate Pol I transcription. Science 373, 547–555 (2021).

63. Hu, S., Lv, P., Yan, Z. & Wen, B. Disruption of nuclear speckles reduces chromatin interactions in active compartments. Epigenetics Chromatin 12, (2019).

64. Park, G. et al. Topology of gene regulatory compartments relative to the nuclear matrix. 2022.05.10.491284 Preprint at https://doi.org/10.1101/2022.05.10.491284 (2022).

65. Guo, Y. E. et al. Pol II phosphorylation regulates a switch between transcriptional and splicing condensates. Nature 572, 543–548 (2019).

66. Lee, D. S. W., Wingreen, N. S. & Brangwynne, C. P. Chromatin mechanics dictates subdiffusion and coarsening dynamics of embedded condensates. Nat. Phys. 17, 531–538 (2021).

67. Gu, B. et al. Transcription-coupled changes in nuclear mobility of mammalian cis-regulatory elements. Science 359, 1050–1055 (2018).

68. Schwalb, B. et al. TT-seq maps the human transient transcriptome. Science 352, 1225–8 (2016).

69. Suter, D. M. et al. Mammalian genes are transcribed with widely different bursting kinetics. 332, 472–474 (2011).

70. Mao, Y. S., Sunwoo, H., Zhang, B. & Spector, D. L. Direct visualization of the co-transcriptional assembly of a nuclear body by noncoding RNAs. Nat. Cell Biol. 13, 95–101 (2011).

71. Pancholi, A. et al. RNA polymerase II clusters form in line with surface condensation on regulatory chromatin. Mol. Syst. Biol. 17, e10272 (2021).

72. Sabari, B. R. et al. Coactivator condensation at super-enhancers links phase separation and gene control. Science 361, (2018).

73. Shrinivas, K. et al. Enhancer features that drive formation of transcriptional condensates. Mol. Cell 75, 549–561.e7 (2019).

74. Markaki, Y. et al. Xist nucleates local protein gradients to propagate silencing across the X chromosome. Cell 0, (2021).

75. Jachowicz, J. W. et al. Xist spatially amplifies SHARP/SPEN recruitment to balance chromosome-wide silencing and specificity to the X chromosome. Nat. Struct. Mol. Biol. 29, 239–249 (2022).

76. Novo, C. L. et al. Satellite repeat transcripts modulate heterochromatin condensates and safeguard chromosome stability in mouse embryonic stem cells. Nat. Commun. 13, 3525 (2022).

77. Gouveia, B. et al. Capillary forces generated by biomolecular condensates. Nature 609, 255–264 (2022).

78. Shin, Y. et al. Spatiotemporal Control of Intracellular Phase Transitions Using Light-Activated optoDroplets. Cell 168, 159–171.e14 (2017).

79. Bracha, D. et al. Mapping Local and Global Liquid Phase Behavior in Living Cells Using Photo-Oligomerizable Seeds. Cell 175, 1467–1480.e13 (2018).

80. Mumford, T. R. et al. Visual detection of submicroscopic protein clusters with a phase-separation-based fluorescent reporter. 2022.07.13.499962 Preprint at https://doi.org/10.1101/2022.07.13.499962 (2022).

81. Jacobs, W. M. Self-Assembly of Biomolecular Condensates with Shared Components. Phys. Rev. Lett. 126, 258101 (2021).

82. Shrinivas, K. & Brenner, M. P. Phase separation in fluids with many interacting components. Proc. Natl. Acad. Sci. 118, (2021).

83. Zwicker, D. & Laan, L. Evolved interactions stabilize many coexisting phases in multicomponent liquids. Proc. Natl. Acad. Sci. 119, e2201250119 (2022).

84. Kroschwald, S. et al. Promiscuous interactions and protein disaggregases determine the material state of stress-inducible RNP granules. eLife 4, e06807 (2015).

85. Jankowsky, E. & Harris, M. E. Specificity and nonspecificity in RNA–protein interactions. Nat. Rev. Mol. Cell Biol. 16, 533–544 (2015).

86. Hohenberg, P. C. & Halperin, B. I. Theory of dynamic critical phenomena. Rev Mod Phys 49, 435–479 (1977).

87. Guyer, J. E., Wheeler, D. & Warren, J. A. FiPy: Partial differential equations with python. Comput. Sci. Eng. 11, 6–15 (2009).

88. Niewidok, B. et al. Single-molecule imaging reveals dynamic biphasic partition of RNA-binding proteins in stress granules. J. Cell Biol. jcb.201709007 (2018) doi:10.1083/jcb.201709007.

89. Coté, A. et al. pre-mRNA spatial distributions suggest that splicing can occur post-transcriptionally. 2020.04.06.028092 Preprint at https://doi.org/10.1101/2020.04.06.028092 (2021).

