## Supplementary material for "Organization and regulation of nuclear condensates by gene activity": SI

for

[#]co-first authors, [$]co-corresponding authors

### Supplementary Information

#### Model definition

In general, active nuclear condensates contain nuclear proteins and RNA species. These species can interact with each other via a plethora of interactions including, but not limited to - interactions mediated by disordered domains [84] specific structured domains such as RBDs [85] , and generic electrostatic interactions [41]. These interactions are captured using a free-energy functional described below:

$$F[\phi_P, \phi_R] = \rho_P(\phi_P - \alpha)^2(\phi_P - \beta)^2 - \chi\phi_P\phi_R + c\phi_P{\phi_R}^2 + \rho_R{\phi_R}^2 + \frac{\kappa}{2}|\phi_P|^2$$

Here, $\phi_P$ and $\phi_R$ are concentrations of the nuclear proteins and the RNAs respectively. The first term is a double-well potential that captures protein-protein interactions which drive phase separation, the next two terms capture RNA-protein interactions that result in a re-entrant phase diagram, the last two terms capture the RNA-RNA repulsion and the surface energy of the protein droplet respectively. The choice of parameters used and the justification of these values and the calculation of the Jacobian matrix is discussed in more detail in [27] .

The model used in this paper to study the dynamics of active nuclear condensates is very similar to the one used in [27]. The total amount of protein is treated as a conserved quantity in the time scales of interested and the protein concentration field is assumed to undergo Model B dynamics[86]:

$$\frac{\partial \phi_P}{\partial t} = M_P \nabla^2 \left(\frac{\delta F}{\delta \phi_P}\right)$$

The dynamics of the protein concentration field is coupled to reaction diffusion dynamics of the RNA species:

$$\frac{\partial \phi_R}{\partial t} = M_R \nabla^2 \phi_R + k_P(x)\phi_P - k_d\phi_R$$

The first term on the right hand sides captures the diffusion of the RNA species in the nucleus and the last term is a first order decay of the RNA species. The second term on the RHs, the RNA production term, captures the spatial clustering of genic hubs through a spatially varying Gaussian function, which can be interpreted to be proportional to the local gene density. The RNA transcription rate is a product of this rate constant with the protein concentration, reflecting the increased rate of transcription that is expected when higher concentrations of catalyzing protein machinery is present.

**Simulations**

The model partial differential equations were numerically simulated using a custom python code available at https://github.com/npradeep96/RNA_localization_final . The code uses the finite volume solver Fipy, developed by the National Institute of Standards and Technology[87] under the hood. All simulations in this paper were done in a 2D circular domain of radius 30 units, with a circular discrete mesh. The spatially discretized PDEs were solved for each incremental time step using the sweep() function in Fipy, with adaptive time stepping to pick smaller or larger time steps depending on the how quickly or slowly the concentration fields change. A grid size of $\Delta r = 0.2$ and a typical time step size on the scale of $\Delta t \sim 0.5$ worked well for the simulations. Simulations were run for a duration of about 15000 time steps, which was sufficient for the system to reach a steady state.

The coexistence concentrations for the protein based on the double well potential are $\alpha = 0.1$ and $\beta = 0.7$. Simulations were done by nucleating a dense seed of protein with a concentration $\phi_P = 0.63$ within a background of dilute protein with a concentration $\phi_P = 0.13$ in the nucleation-and-growth regime. The initial concentration of the RNA species is $\phi_R = 0$ everywhere. The no-flux Neumann boundary condition was applied to all species at the domain boundaries.

For simulations varying the width of the gene-dense region RNA-producing region $\sigma$, the integral of the rate constant i.e. $k_T$ was fixed. Since the rate constant is given by the Gaussian expression $k_P(x) = ce^{-||x-x_0||^2/2\sigma^2}$, the constant $c$ is chosen for each value of sigma such that the integral $k_T = \int_0^R \; k_p(x).2\pi x.dx = constant.$ An approximate value of the constant given $k_T$ and $\sigma$ is $c \approx \frac{k_T}{\int_0^\infty \; e^{-||x-x_0||^2/2\sigma^2}.2\pi x.dx} = \frac{k_T}{2\pi\sigma^2}$.

**Analyses**

To calculate the radii of the vacuoles if Figure 3, we calculated the radius of the region of dilute protein concentration ($\phi_P < (\frac{\alpha+\beta}{2} = 0.4)$) within the dense condensate of protein with ($\phi_P > 0.4$). The reaction-diffusion length scale for Figure 4, $l_{r-d}$ was calculated as $l_{r-d} = \frac{M_R}{k_{P,avg}}$ where $k_{P,avg} = \frac{\int k_P(x).2\pi x.dx}{\int 2\pi x.dx}$ The centroid of the protein concentration profiles in Figure 6 was calculated as the center of mass of this concentration profile $\phi_p(x)$. The eccentricity was calculated as the $e = \frac{(I_{xx}-I_{yy})^2-4I_{xy}}{(I_{xx}+I_{yy})^2}$, where $I_{xx}$ and $I_{yy}$ are the second moments of the concentration profile along the x and the y directions, and $I_{xy}$ is the cross moment.

**Gradient calculation**

RNAs in the nucleus exhibit a range of diffusivities, with typically chromatin-associated mobilities around ~ $10^{-3.5}\mu m^2/s$ but higher for transported for mRNPs [68,88]. Depending on the type of RNA species expressed, median half-lives range from ~1 min (for nc and eRNAs) to 50 mins (for lnc and mRNAs) [69] and mammalian RNAs are transcribed across a wide range of rates, with some typical ranges for mRNAs between~ 0.5 - 3 mRNAs/min [70]. Since its not apriori clear whether the *on* or *off* rates are limiting *in vivo* (by contrast, in our models, $k_d$ is high so the gradient length-scale is set largely by $< k_p >$). Hence we estimate an approximate range of gradient length-scales as $g_{off} = \sqrt{M_{rna}/k_{off}} \approx 0.2 - 1\ \mu m$ (assuming similar on-rates) and $g_{on} = \sqrt{M_{rna}/k_{on}} \approx 0.08 - 0.25\ \mu m$ (assuming fast and similar koff rates). These length scales are broadly consistent with the observations that transcribing and non-coding RNAs are often proximal, within a micron or so, to their site of transcription [40,89]

#### Parameters for simulations

**Figure 2A & 2B:**

| | |
|---|---|
| Free energy | $\alpha = 0.1, \beta = 0.7, \rho_P = 1.0, \chi = 1.0, c = 10.0, \rho_R = 10.0$ and $\kappa = 0.5$ |
| Initial conditions | Nucleated dense phase of protein at (0,0): $r = 4.0, \phi_P = 0.63$<br>Background protein concentration: $\phi_P = 0.12$ |
| Kinetic parameters | $M_P = 1.0, M_R = 1.0, k_d = 0.5, \sigma = 2.0$ |
| Numerical integration | $\Delta t_{min} = 1 \times 10^{-8}, \Delta t_{max} = 0.5, total\ steps = 15000$ |
| Geometry | 2D circular domain: $r_{domain} = 30$. Mesh size: $\Delta r = 0.4$. No flux boundary condition for all species. |
| Parameters varied | $k_T = [0.1\ to\ 100]$ (for the case with gene compartment)<br>$k_T = [0.1\ to\ 1413]$ (for the uniform case) |

**Figure 2C & 2D:**

| | |
|---|---|
| Free energy | $\alpha = 0.1, \beta = 0.7, \rho_P = 1.0, \chi = 1.0, c = 10.0, \rho_R = 10.0$ and $\kappa = 0.5$ |
| Initial conditions | Nucleated dense phase of protein at (0,0): $r = 4.0, \phi_P = 0.63$<br>Background protein concentration: $\phi_P = 0.12$ |
| Kinetic parameters | $M_P = 1.0, M_R = 1.0, k_d = 0.5, k_T = 25\ or\ 50$ |
| Numerical integration | $\Delta t_{min} = 1 \times 10^{-8}, \Delta t_{max} = 0.5, total\ steps = 15000$ |
| Geometry | 2D circular domain: $r_{domain} = 30$. Mesh size: $\Delta r = 0.4$. No flux boundary |

| | |
|---|---|
| | condition for all species. |
| Parameters varied | $\sigma = [1\ to\ 8]$ |

**Figure 2E:**

| | |
|---|---|
| Free energy | $\alpha = 0.1, \beta = 0.7, \rho_P = 1.0, \chi = 1.0, c = 10.0, \rho_R = 10.0$ and $\kappa = 0.5$ |
| Initial conditions | No nucleated dense phase of protein at (0,0).<br>Background protein concentration: $\phi_P = 0.12$ |
| Kinetic parameters | $M_P = 1.0, M_R = 1.0, k_d = 0.5, \sigma = 7.5$ |
| Numerical integration | $\Delta t_{min} = 1 \times 10^{-8}, \Delta t_{max} = 0.5, total\ steps = 15000$ |
| Geometry | 2D circular domain: $r_{domain} = 30$. Mesh size: $\Delta r = 0.4$. No flux boundary condition for all species. |
| Parameters varied | $k_T = [1\ to\ 200]$ |

**Figure 2F:**

| | |
|---|---|
| Free energy | $\alpha = 0.1, \beta = 0.7, \rho_P = 1.0, \chi = 1.0, c = 10.0, \rho_R = 10.0$ and $\kappa = 0.5$ |
| Initial conditions | No nucleated dense phase of protein at (0,0).<br>Background protein concentration: $\phi_P = 0.12$ |
| Kinetic parameters | $M_P = 1.0, M_R = 1.0, k_d = 0.5$ |
| Numerical integration | $\Delta t_{min} = 1 \times 10^{-8}, \Delta t_{max} = 0.5, total\ steps = 15000$ |

| | |
|---|---|
| Geometry | 2D circular domain: $r_{domain} = 30$. Mesh size: $\Delta r = 0.4$. No flux boundary condition for all species. |
| Parameters varied | $k_T = [1\ to\ 200], \sigma = [1\ to\ 8]$ |

**Figure 3A:**

| | |
|---|---|
| Free energy | $\alpha = 0.1, \beta = 0.7, \rho_P = 1.0, \chi = 1.0, c = 10.0, \rho_R = 10.0$ and $\kappa = 0.5$ |
| Initial conditions | Nucleated dense phase of protein at (0,0): $r = 4.0, \phi_P = 0.63$<br>Background protein concentration: $\phi_P = 0.13$ |
| Kinetic parameters | $M_P = 1.0, M_R = 1.0, k_d = 0.5, \sigma = 3$ |
| Numerical integration | $\Delta t_{min} = 1 \times 10^{-8}, \Delta t_{max} = 0.5, total\ steps = 15000$ |
| Geometry | 2D circular domain: $r_{domain} = 30$. Mesh size: $\Delta r = 0.2$. No flux boundary condition for all species. |
| Parameters varied | $k_T = 1.0$ (top panel)<br>$k_T = 20.0$ (bottom panel) |

**Figure 3B:**

| | |
|---|---|
| Free energy | $\alpha = 0.1, \beta = 0.7, \rho_P = 1.0, \chi = 1.0, c = 10.0, \rho_R = 10.0$ and $\kappa = 0.5$ |
| Initial conditions | Nucleated dense phase of protein at (0,0): $r = 4.0, \phi_P = 0.63$ |

| | |
|---|---|
| | Background protein concentration: $\phi_P = 0.13$ |
| Kinetic parameters | $M_P = 1.0, M_R = 1.0, k_d = 0.5$ |
| Numerical integration | $\Delta t_{min} = 1 \times 10^{-8}, \Delta t_{max} = 0.5, total\ steps = 15000$ |
| Geometry | 2D circular domain: $r_{domain} = 30$. Mesh size: $\Delta r = 0.2$. No flux boundary condition for all species. |
| Parameters varied | $k_T = [1\ to\ 200]$, $\sigma = 4$ (Left panel)<br>$\sigma = [2\ to\ 8], k_T = 90.0$ (Right panel) |

**Figure 3C:**

| | |
|---|---|
| Free energy | $\alpha = 0.1, \beta = 0.7, \rho_P = 1.0, \chi = 1.0, c = 10.0, \rho_R = 10.0$ and $\kappa = 0.5$ |
| Initial conditions | Nucleated dense phase of protein at (0,0): $r = 4.0, \phi_P = 0.63$<br>Background protein concentration: $\phi_P = 0.13$ |
| Kinetic parameters | $M_P = 1.0, M_R = 1.0, k_d = 0.5, \sigma = 4, k_T = 50$ |
| Numerical integration | $\Delta t_{min} = 1 \times 10^{-8}, \Delta t_{max} = 0.5, total\ steps = 15000$ |
| Geometry | 2D circular domain: $r_{domain} = 30$. Mesh size: $\Delta r = 0.2$. No flux boundary condition for all species. |
| Parameters varied | For an initial protein concentration of $\phi_{P0}$ at each position in space, a Gaussian noise with variance 0.1$\phi_{P0}$ and zero mean was added, which resulted in the vacuoles exhibiting shape instabilities and aspherical morphologies with different centroids. |

**Figure 4A:**

| | |
|---|---|
| Free energy | $\alpha = 0.1, \beta = 0.7, \rho_P = 1.0, \chi = 1.0, c = 10.0, \rho_R = 10.0$ and $\kappa = 0.5$ |
| Initial conditions | Nucleated dense phase of protein at (0,0): $r = 4.0, \phi_P = 0.63$<br>Background protein concentration: $\phi_P = 0.12$ |
| Kinetic parameters | $M_P = 1.0, M_R = 1.0, k_d = 0.5, k_T = 10, \sigma = 5$ |
| Numerical integration | $\Delta t_{min} = 1 \times 10^{-8}, \Delta t_{max} = 0.5, total\ steps = 15000$ |
| Geometry | 2D circular domain: $r_{domain} = 30$. Mesh size: $\Delta r = 0.2$. No flux boundary condition for all species. |
| Parameters varied | The center of the gene activity is located at distance $r = 10$ from the nucleated dense phase of protein. |

**Figure 4B:**

| | |
|---|---|
| Free energy | $\alpha = 0.1, \beta = 0.7, \rho_P = 1.0, \chi = 1.0, c = 10.0, \rho_R = 10.0$ and $\kappa = 0.5$ |
| Initial conditions | Nucleated dense phase of protein at (0,0): $r = 4.0, \phi_P = 0.63$<br>Background protein concentration: $\phi_P = 0.13$ |
| Kinetic parameters | $M_P = 1.0, M_R = 1.0, k_d = 0.5, k_T = 10, \sigma = 4$ |
| Numerical integration | $\Delta t_{min} = 1 \times 10^{-8}, \Delta t_{max} = 0.5, total\ steps = 15000$ |
| Geometry | 2D circular domain: $r_{domain} = 30$. Mesh size: $\Delta r = 0.2$. No flux boundary condition for all species. |
| Parameters varied | The center of the gene activity is located at distance $r = [2\ to\ 25]$ from the |

| | |
|---|---|
| | nucleated dense phase of protein. |

**Figure 4C:**

| | |
|---|---|
| Free energy | $\alpha = 0.1, \beta = 0.7, \rho_P = 1.0, \chi = 1.0, c = 10.0, \rho_R = 10.0$ and $\kappa = 0.5$ |
| Initial conditions | Nucleated dense phase of protein at (0,0): $r = 4.0, \phi_P = 0.63$<br>Background protein concentration: $\phi_P = 0.13$<br>The center of the gene activity is located at distance $r = [15]$ from the nucleated dense phase of protein. |
| Kinetic parameters | $M_P = 1.0, k_d = 0.5, k_T = 10, \sigma = 4$ |
| Numerical integration | $\Delta t_{min} = 1 \times 10^{-8}, \Delta t_{max} = 0.5, total\ steps = 15000$ |
| Geometry | 2D circular domain: $r_{domain} = 30$. Mesh size: $\Delta r = 0.2$. No flux boundary condition for all species. |
| Parameters varied | $M_R = [10^{-4}\ to\ 200]$ |

**Figure 5:**

| | |
|---|---|
| Free energy | $\alpha = 0.1, \beta = 0.7, \rho_P = 1.0, \chi = 1.0, c = 10.0, \rho_R = 10.0$ and $\kappa = 0.5$ |
| Initial conditions | Nucleated dense phase of protein at (0,0): $r = 4.0, \phi_P = 0.63$<br>Background protein concentration: $\phi_P = 0.13$ |
| Kinetic parameters | $M_P = 1.0, M_R = 1.0, k_d = 0.5, \sigma = 4$ |
| Numerical integration | $\Delta t_{min} = 1 \times 10^{-8}, \Delta t_{max} = 0.5, total\ steps = 15000$ |

| | |
|---|---|
| Geometry | 2D circular domain: $r_{domain} = 30$. Mesh size: $\Delta r = 0.2$. No flux boundary condition for all species. |
| Parameters varied | $k_T = [1\ to\ 200]$<br>The center of the gene activity is located at distances $r = [0\ to\ 25]$ from the nucleated dense phase of protein. |

**Figure 6A:**

| | |
|---|---|
| Free energy | $\alpha = 0.1, \beta = 0.7, \rho_P = 1.0, \chi = 1.0, c = 10.0, \rho_R = 10.0$ and $\kappa = 0.5$ |
| Initial conditions | Nucleated dense phase of protein at (0,0): $r = 4.0, \phi_P = 0.63$<br>Background protein concentration: $\phi_P = 0.13$<br>The distance between the sites of gene A and gene B is $r = 10$. The nucleated condensate is located at the mid point of the line joining the sites of activity. |
| Kinetic parameters | $M_P = 1.0, M_R = 1.0, k_d = 0.5, \sigma_A = \sigma_B = 4$ |
| Numerical integration | $\Delta t_{min} = 1 \times 10^{-8}, \Delta t_{max} = 0.5, total\ steps = 15000$ |
| Geometry | 2D circular domain: $r_{domain} = 30$. Mesh size: $\Delta r = 0.4$. No flux boundary condition for all species. |
| Parameters varied | $k_{TB} = 1, k_{TA} = [10^{-4}\ to\ 100]$ |

**Figure 6B:**

| | |
|---|---|
| Free energy | $\alpha = 0.1, \beta = 0.7, \rho_P = 1.0, \chi = 1.0, c = 10.0, \rho_R = 10.0$ and $\kappa = 0.5$ |
| Initial conditions | Nucleated dense phase of protein at (0,0): $r = 4.0, \phi_P = 0.63$<br>Background protein concentration: $\phi_P = 0.13$<br>The distance between the sites of gene A and gene B is $r = 10$. The nucleated condensate is located at the mid point of the line joining the sites of activity. |
| Kinetic parameters | $M_P = 1.0, M_R = 1.0, k_d = 0.5, k_{TA} = k_{TB} = 10$ |
| Numerical integration | $\Delta t_{min} = 1 \times 10^{-8}, \Delta t_{max} = 0.5, total\ steps = 15000$ |
| Geometry | 2D circular domain: $r_{domain} = 30$. Mesh size: $\Delta r = 0.2$. No flux boundary condition for all species. |
| Parameters varied | $\sigma_A = 4, \sigma_B = [1\ to\ 10]$ |

**Figure 6C:**

| | |
|---|---|
| Free energy | $\alpha = 0.1, \beta = 0.7, \rho_P = 1.0, \chi = 1.0, c = 10.0, \rho_R = 10.0$ and $\kappa = 0.5$ |
| Initial conditions | Nucleated dense phase of protein at (0,0): $r = 4.0, \phi_P = 0.63$<br>Background protein concentration: $\phi_P = 0.13$ |
| Kinetic parameters | $M_P = 1.0, M_R = 1.0, k_d = 0.5, k_{TA} = k_{TB} = 10, \sigma_A = \sigma_B = 4$ |
| Numerical integration | $\Delta t_{min} = 1 \times 10^{-8}, \Delta t_{max} = 0.5, total\ steps = 15000$ |

| | |
|---|---|
| Geometry | 2D circular domain: $r_{domain} = 30$. Mesh size: $\Delta r = 0.4$. No flux boundary condition for all species. |
| Parameters varied | The distance between the sites of gene A and gene B is $r = [10\ to\ 24]$. The nucleated condensate is located at the mid point of the line joining the sites of activity. |

**Figure S1B:**

| | |
|---|---|
| Free energy | $\alpha = 0.1, \beta = 0.7, \rho_P = 1.0, \chi = 1.0, c = 10.0, \rho_R = 10.0$ and $\kappa = 0.5$ |
| Initial conditions | Nucleated dense phase of protein at (0,0): $r = 4.0, \phi_P = 0.63$<br>Background protein concentration: $\phi_P = 0.12$ |
| Kinetic parameters | $M_P = 1.0, M_R = 1.0, k_d = 0.5, \sigma = 2.0$ |
| Numerical integration | $\Delta t_{min} = 1 \times 10^{-8}, \Delta t_{max} = 0.5, total\ steps = 15000$ |
| Geometry | 2D circular domain: $r_{domain} = 30$. Mesh size: $\Delta r = 0.4$. No flux boundary condition for all species. |
| Parameters varied | $k_T = [0.5, 0.8, 1.0, 10.0, 25.0]$ (for the case with gene compartment) |

**Figure S1C:**

| | |
|---|---|
| Free energy | $\alpha = 0.1, \beta = 0.7, \rho_P = 1.0, \chi = 1.0, c = 10.0, \rho_R = 10.0$ and $\kappa = 0.5$ |
| Initial conditions | Nucleated dense phase of protein at (0,0): $r = 4.0, \phi_P = 0.63$<br>Background protein concentration: $\phi_P = 0.12$ |

| | |
|---|---|
| Kinetic parameters | $M_P = 1.0, M_R = 1.0, k_d = 0.5$ |
| Numerical integration | $\Delta t_{min} = 1 \times 10^{-8}, \Delta t_{max} = 0.5,\ total\ steps = 15000$ |
| Geometry | 2D circular domain: $r_{domain} = 30$. Mesh size: $\Delta r = 0.4$. No flux boundary condition for all species. |
| Parameters varied | For exponential distribution, $k_p(x) = ce^{-\gamma x}$, with $\gamma = 1\ and\ 4$<br>Fior Gaussian distribution, $k_p(x) = ce^{-x^2/2\sigma^2}$ with $\sigma = 2.0$ |

**Figure S1D:**

| | |
|---|---|
| Free energy | $\alpha = 0.1, \beta = 0.7, \rho_P = 1.0, \chi = 1.0, c = 10.0, \rho_R = 10.0$ and $\kappa = 0.5$ |
| Initial conditions | No nucleated dense phase of protein at (0,0).<br>Background protein concentration: $\phi_P = 0.12$ |
| Kinetic parameters | $M_P = 1.0, M_R = 1.0, k_d = 0.5$ |
| Numerical integration | $\Delta t_{min} = 1 \times 10^{-8}, \Delta t_{max} = 0.5,\ total\ steps = 15000$ |
| Geometry | 2D circular domain: $r_{domain} = 30$. Mesh size: $\Delta r = 0.4$. No flux boundary condition for all species. |
| Parameters varied | $k_T = [1\ to\ 200], \sigma = [1\ to\ 8]$ |

**Figure S1E:**

| | |
|---|---|
| Free energy | $\alpha = 0.1, \beta = 0.7, \rho_P = 1.0, \chi = 1.0, c = 10.0, \rho_R = 10.0$ and $\kappa = 0.5$ |

| | |
|---|---|
| Initial conditions | No nucleated dense phase of protein at (0,0).<br>Background protein concentration: $\phi_P = 0.12$ |
| Kinetic parameters | $M_P = 1.0, M_R = 1.0, k_d = 0.5, \sigma = 7.5$ |
| Numerical integration | $\Delta t_{min} = 1 \times 10^{-8}, \Delta t_{max} = 0.5, total\ steps = 15000$ |
| Geometry | 2D circular domain: $r_{domain} = 30$. Mesh size: $\Delta r = 0.4$. No flux boundary condition for all species. |
| Parameters varied | $k_T = [1\ to\ 200]$ |

**Figure S2A:**

| | |
|---|---|
| Free energy | $\alpha = 0.1, \beta = 0.7, \rho_P = 1.0, \chi = 1.0, c = 10.0, \rho_R = 10.0$ and $\kappa = 0.5$ |
| Initial conditions | Nucleated dense phase of protein at (0,0): $r = 4.0, \phi_P = 0.63$<br>Background protein concentration: $\phi_P = 0.13$ |
| Kinetic parameters | $M_P = 1.0, M_R = 1.0, k_d = 0.5$ |
| Numerical integration | $\Delta t_{min} = 1 \times 10^{-8}, \Delta t_{max} = 0.5, total\ steps = 15000$ |
| Geometry | 2D circular domain: $r_{domain} = 30$. Mesh size: $\Delta r = 0.2$. No flux boundary condition for all species. |
| Parameters varied | $k_T = [1\ to\ 200]$, $\sigma = 4$ (Left panel)<br>$\sigma = [2\ to\ 8], k_T = 90.0$ (Right panel) |

**Figure S2B and S2C:**

| | |
|---|---|
| Free energy | $\alpha = 0.1, \beta = 0.7, \rho_P = 1.0, \chi = 1.0, c = 10.0, \rho_R = 10.0$ and $\kappa = 0.5$ |
| Initial conditions | Nucleated dense phase of protein at (0,0): $r = 4.0, \phi_P = 0.63$<br>Background protein concentration: $\phi_P = 0.13$ |
| Kinetic parameters | $M_P = 1.0, M_R = 1.0, k_d = 0.5, \sigma = 4, k_T = 50$ |
| Numerical integration | $\Delta t_{min} = 1 \times 10^{-8}, \Delta t_{max} = 0.5, total\ steps = 15000$ |
| Geometry | 2D circular domain: $r_{domain} = 30$. Mesh size: $\Delta r = 0.2$. No flux boundary condition for all species. |
| Parameters varied | For an initial protein concentration of $\phi_{P0}$ at each position in space, a Gaussian noise with variance 0.1$\phi_{P0}$ and zero mean was added, which resulted in the vacuoles exhibiting shape instabilities and aspherical morphologies with different centroids. |

**Figure 2D:**

| | |
|---|---|
| Free energy | $\alpha = 0.1, \beta = 0.7, \rho_P = 1.0, \chi = 1.0, c = 10.0, \rho_R = 10.0$ and $\kappa = 0.5$ |
| Initial conditions | Nucleated dense phase of protein at (0,0): $r = 4.0, \phi_P = 0.63$<br>Background protein concentration: $\phi_P = 0.13$ |
| Kinetic parameters | $M_P = 1.0, M_R = 1.0, k_d = 0.5, \sigma = 3, k_T = 50$ |
| Numerical integration | $\Delta t_{min} = 1 \times 10^{-8}, \Delta t_{max} = 0.5, total\ steps = 15000$ |
| Geometry | 2D circular domain: $r_{domain} = 30$. Mesh size: $\Delta r = 0.2$. No flux boundary condition for all species. |
| Parameters varied | $\kappa = [0.5\ to\ 2.0]$ (for $k_T = 30$) |

| | |
|---|---|
| | $\kappa = [0.01\ to\ 0.4]$ (for $k_T = 50$) |

**Figure S2E:**

| | |
|---|---|
| Free energy | $\alpha = 0.1, \beta = 0.7, \rho_P = 1.0, \chi = 1.0, c = 10.0, \rho_R = 10.0$ and $\kappa = 0.5$ |
| Initial conditions | Nucleated dense phase of protein at (0,0): $r = 4.0, \phi_P = 0.63$<br>Background protein concentration: $\phi_P = 0.13$ |
| Kinetic parameters | $M_P = 1.0, M_R = 1.0, k_d = 0.5, \sigma = 2, k_T = 50$ |
| Numerical integration | $\Delta t_{min} = 1 \times 10^{-8}, \Delta t_{max} = 0.5, total\ steps = 15000$ |
| Geometry | 2D circular domain: $r_{domain} = 30$. Mesh size: $\Delta r = 0.2$. No flux boundary condition for all species. |
| Parameters varied | $M_R = [0.1\ to\ 10]$ |

**Figure S3A:**

| | |
|---|---|
| Free energy | $\alpha = 0.1, \beta = 0.7, \rho_P = 1.0, \chi = 1.0, c = 10.0, \rho_R = 10.0$ and $\kappa = 0.5$ |
| Initial conditions | Nucleated dense phase of protein at (0,0): $r = 4.0, \phi_P = 0.63$<br>Background protein concentration: $\phi_P = 0.13$<br>The center of the gene activity is located at distance $r = [15]$ from the nucleated dense phase of protein. |
| Kinetic parameters | $M_P = 1.0, k_d = 0.5, k_T = 10, \sigma = 4$ |
| Numerical integration | $\Delta t_{min} = 1 \times 10^{-8}, \Delta t_{max} = 0.5, total\ steps = 15000$ |

| | |
|---|---|
| Geometry | 2D circular domain: $r_{domain} = 30$. Mesh size: $\Delta r = 0.2$. No flux boundary condition for all species. |
| Parameters varied | $M_R = [10 \quad to\ 200]$ |

**Figure S3B:**

| | |
|---|---|
| Free energy | $\alpha = 0.1, \beta = 0.7, \rho_P = 1.0, \chi = 1.0, c = 10.0, \rho_R = 10.0$ |
| Initial conditions | Nucleated dense phase of protein at (0,0): $r = 4.0, \phi_P = 0.63$<br>Background protein concentration: $\phi_P = 0.13$<br>The center of the gene activity is located at distance $r = [15]$ from the nucleated dense phase of protein. |
| Kinetic parameters | $M_P = 1.0, M_R = 1.0, \quad k_d = 0.5, k_T = 10, \sigma = 4$ |
| Numerical integration | $\Delta t_{min} = 1 \times 10^{-8}, \Delta t_{max} = 0.5, total\ steps = 15000$ |
| Geometry | 2D circular domain: $r_{domain} = 30$. Mesh size: $\Delta r = 0.2$. No flux boundary condition for all species. |
| Parameters varied | $\kappa = [0.01\ to\ 0.5]$ |

**Figure S4A**

| | |
|---|---|
| Free energy | $\alpha = 0.1, \beta = 0.7, \rho_P = 1.0, \chi = 1.0, c = 10.0, \rho_R = 10.0$ |
| Initial conditions | Nucleated dense phase of protein at (0,0): $r = 4.0, \phi_P = 0.63$<br>Background protein concentration: $\phi_P = 0.13$ |

| | |
|---|---|
| Kinetic parameters | $M_P = 1.0, M_R = 1.0, \quad k_d = 0.5, k_{TA} = k_{TB} = 1, \sigma_A = \sigma_B = 4$ |
| Numerical integration | $\Delta t_{min} = 1 \times 10^{-8}, \Delta t_{max} = 0.5, total\ steps = 15000$ |
| Geometry | 2D circular domain: $r_{domain} = 30$. Mesh size: $\Delta r = 0.2$. No flux boundary condition for all species. |
| Parameters varied | $\kappa = [0.005\ to\ 0.5]$<br>The distance between the centers of gene A and gene B was varied from $r/2\sigma = [0\ to\ 2.5]$ |

**Figure S4B & S4C**

| | |
|---|---|
| Free energy | $\alpha = 0.1, \beta = 0.7, \rho_P = 1.0, \chi = 1.0, c = 10.0, \rho_R = 10.0, \kappa = 0.05$ |
| Initial conditions | Nucleated dense phase of protein at (0,0): $r = 4.0, \phi_P = 0.63$<br>Background protein concentration: $\phi_P = 0.13$ |
| Kinetic parameters | $M_P = 1.0, M_R = 1.0, \quad k_d = 0.5, k_{TA} = k_{TB} = 1, \sigma_A = \sigma_B = 4$ |
| Numerical integration | $\Delta t_{min} = 1 \times 10^{-8}, \Delta t_{max} = 0.5, total\ steps = 15000$ |
| Geometry | 2D circular domain: $r_{domain} = 30$. Mesh size: $\Delta r = 0.2$. No flux boundary condition for all species. |
| Parameters varied | The distance between the centers of gene A and gene B was varied from $r/2\sigma = [1, 1.5, 2, 2.5]$ |

**Figure S4D**

| | |
|---|---|
| Free energy | $\alpha = 0.1, \beta = 0.7, \rho_P = 1.0, \chi = 1.0, c = 10.0, \rho_R = 10.0, \kappa = 0.05$ |
| Initial conditions | Nucleated dense phase of protein at (0,0): $r = 4.0, \phi_P = 0.63$<br>Background protein concentration: $\phi_P = 0.13$ |
| Kinetic parameters | $M_P = 1.0, M_R = 1.0, \quad k_d = 0.5, k_{TA} = k_{TB} = 1, \sigma_A = \sigma_B = 4$ |
| Numerical integration | $\Delta t_{min} = 1 \times 10^{-8}, \Delta t_{max} = 0.5, total\ steps = 15000$ |
| Geometry | 2D circular domain: $r_{domain} = 30$. Mesh size: $\Delta r = 0.2$. No flux boundary condition for all species.<br>The distance between the centers of gene A and gene B was fixed at $r/2\sigma = 2.5$ |
| Parameters varied | $\kappa = [0.01, 0.05, 0.1, 0.5]$ |

#### Supplementary Figure Captions

**Figure S1. Effect of different parameters of RNA activity and free-energy on condensate size and nucleation**.

A. Dynamic evolution of radius of condensate for select kT values over time.
B. Effect of selected distribution of kT. Plotted are steady-state condensate radius as a function of $k_T$ where $k_P(x)$ is either an exponential distribution with γ = 1 or γ = 4, or a gaussian distribution, which is used for the bulk of this study.
C. Steady-state condensate radius (left-panel) vs total RNA activity (kT) for different spread in RNA activity (σ). In general, increasing spread leads to decreasing local RNA concentrations and larger stable condensates.
D. Steady-state radius (left) and characteristic nucleation time (right) as a function of $k_T$ in conditions with no initial condensate for spatially clustered and uniform RNA production.

**Figure S2. Non-equilibrium steady-state morphologies are modulated by surface tension and RNA mobility**

A. Variance of vacuole size (color of dot) versus total gene activity or extent of clustering.
B. Histogram of centroids of aspherical droplets formed after symmetry-breaking for different initial conditions.
C. In few simulations, symmetry-breaking initially leads to two aspherical condensates that eventually fuse into one at steady-state.
D. Role of surface tension in symmetry breakage and vacuole formation. Left: at kT= 30 with kT(x) modeled by a Gaussian distribution with std = 4, increasing surface tension causes a vacuole to break symmetry and eventually dissolve. Right: kT= 50 with kT(x) modeled by a Gaussian distribution with std = 4, decreasing surface tension causes a condensate which initially broke symmetry to stabilize and form a vacuole.
E. The effect of RNA mobility on condensate morphologies at steady state. Left: condensate radii at steady state as a function of RNA mobility coefficient.

**Figure S3. Flow of condensates is characterized by surface tension and strength of RNA gradient**

A. Speed of condensate in regimes where flow is observed as a function of RNA mobility coefficient. Top: dynamic velocity of condensate over time. Bottom: peak velocity of condensate as a function of RNA mobility coefficient.
B. Condensate behavior in response to changes in surface tension within regimes where flow is observed (r = 10, kT = 10.0). Top: distance between initial condensate and center of RNA source as a function of time. Bottom: eccentricity of condensate as a function of time.

**Figure S4. Two sites of activity can lead to droplet division**

A. Phase diagram for condensate morphology upon varying the surface tension of the protein ($\kappa$) and the distance between two active genomic loci $r$. The distance between the loci is reported as a dimensionless ratio with the width of the genomic locus ($r/2\sigma$). In the parameter regime 1 - we have a single droplet at steady state. In parameter regime 2 where the surface tension is low and the loci are far apart, the system at steady state has two droplets each centered around one of the loci.
B. Dynamics of the eccentricity of the droplets for different distances between the loci ($r/2\sigma$). As the distance between the loci increases, the eccentricity of the droplet increases more rapidly as the droplet reaches a deformed steady state shape. Beyond a threshold distance (light grey), the single droplet deforms so much after a certain time that it quickly splits into two spherical droplets which is a more energetically favorable configuration.
C. Steady state droplet morphology upon varying distance between the loci. Increasing the distance between the loci leads to a gradual deformation of the droplet. Beyond a threshold distance, the droplet splits into two, each one centered around one of the loci.
D. Steady state droplet morphology upon varying the surface tension ($\kappa$). Increasing the surface tension penalizes interface formation and progressively favors a single droplet configuration.
