## Supplementary material for "Organization and regulation of nuclear condensates by gene activity": SI_figures

**Figure S1.  Effect of different parameters of RNA activity and free-energy on condensate size and nucleation.**

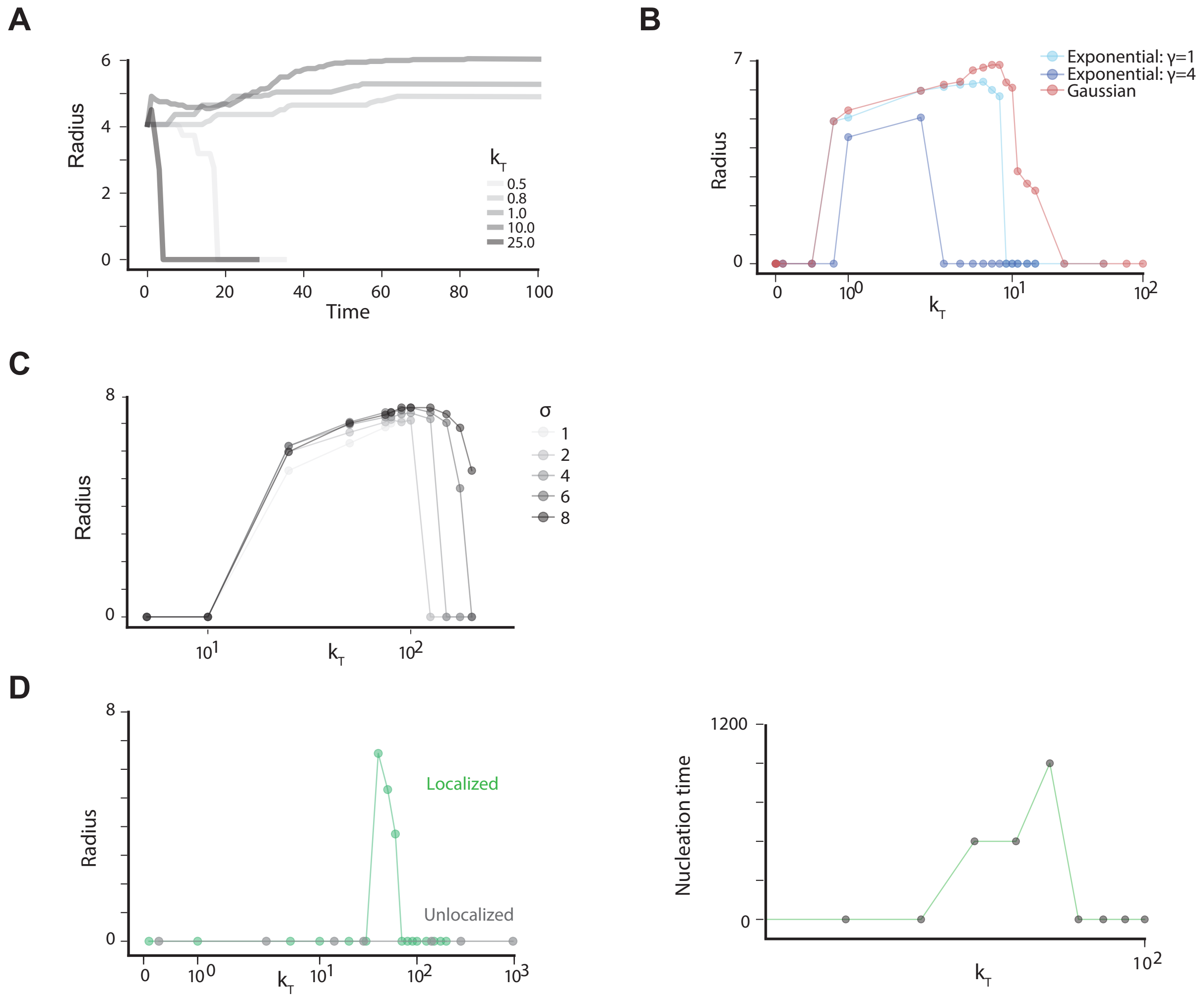


**Figure S1.  Effect of different parameters of RNA activity and free-energy on condensate size and nucleation.**

A. Dynamic evolution of radius of condensate for select $k_T$ values over time.

B. Effect of selected distribution of $k_T$. Plotted are steady-state condensate radius as a function of $k_T$ where $k_P(x)$ is either an exponential distribution with γ = 1 or γ = 4, or a gaussian distribution, which is used for the bulk of this study.

C. Steady-state condensate radius (left-panel) vs total RNA activity ($k_T$) for different spread in RNA activity (σ). In general,increasing spread leads to decreasing local RNA concentrations and larger stable condensates.

D. Steady-state radius (left) and characteristic nucleation time (right) as a function of $k_T$ in conditions with no initial condensate for spatially clustered and uniform RNA production.

**Figure S2. Non-equilibrium steady-state morphologies are modulated by surface tension and RNA mobility**

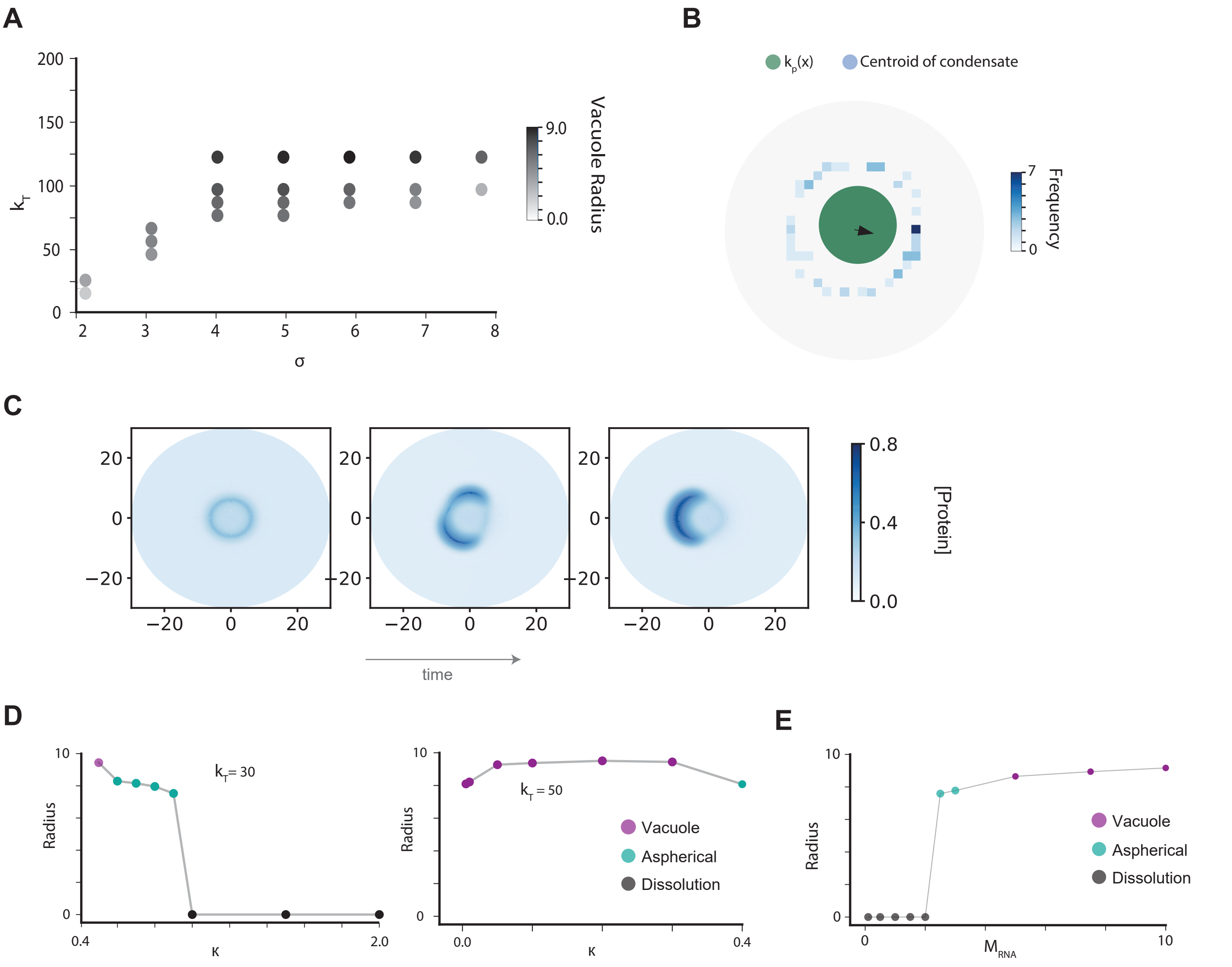


**Figure S2. Non-equilibrium steady-state morphologies are modulated by surface tension and RNA mobility**

A. Variance of vacuole size (color of dot) versus total gene activity or extent of clustering.

B. Histogram of centroids of aspherical droplets formed after symmetry-breaking for different initial conditions.

C. In few simulations, symmetry-breaking initially leads to two aspherical condensates that eventually fuse into one at steady-state.

D. Role of surface tension in symmetry breakage and vacuole formation. Left: at $k_T$= 30 with $k_T(x)$ modelled by a Gaussian distribution with std = 4, increasing surface tension causes a vacuole to break symmetry and eventually dissolve. Right: $k_T$= 50 with $k_T(x)$ modelled by a Gaussian distribution with std = 4, decreasing surface tension causes a condensate which initially broke symmetry to stabilize and form a vacuole.

E. The effect of RNA mobility on condensate morphologies at steady state. Left: condensate radii at steady state as a function of RNA mobility coefficient. Increasing mobility decreases local RNA concentration, which leads to condensate stabilization, first in an aspherical droplet, eventually to a symmetric vacuole.

**Figure S3. Flow of condensates is characterized by surface tension and strength of RNA gradient**

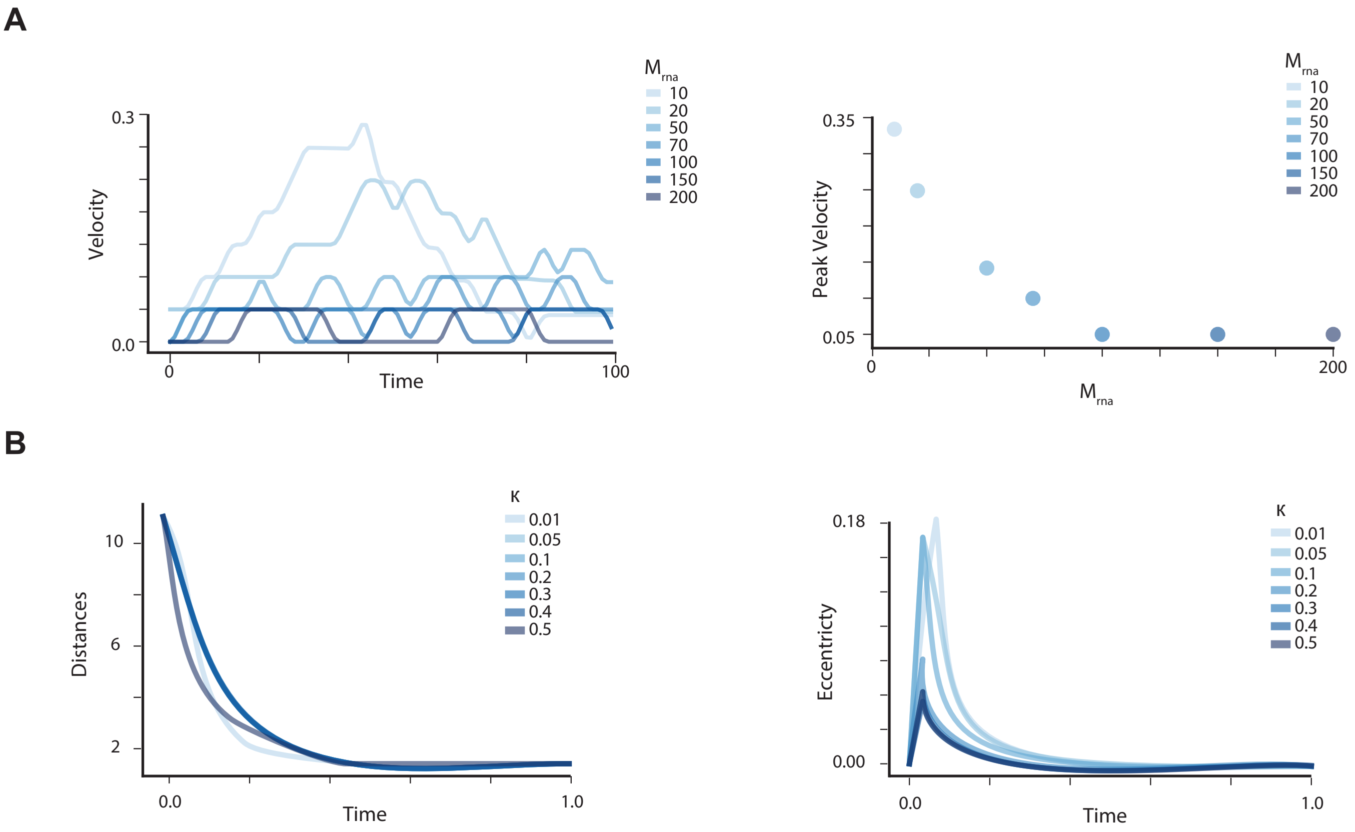


**Figure S3. Flow of condensates is characterized by surface tension and strength of RNA gradient**

A. Speed of condensate in regimes where flow is observed as a function of RNA mobility coefficient. Top: dynamic velocity of condensate over time. Bottom: peak velocity of condensate as a function of RNA mobility coefficient.

B. Condensate behavior in response to changes in surface tension within regimes where flow is observed (r = 10, $k_T$ = 10.0). Top: distance between initial condensate and center of RNA source as a function of time. Bottom: eccentricity of condensate as a function of time.

**Figure S4. Two sites of RNA activity can lead to droplet division and other morphologies**

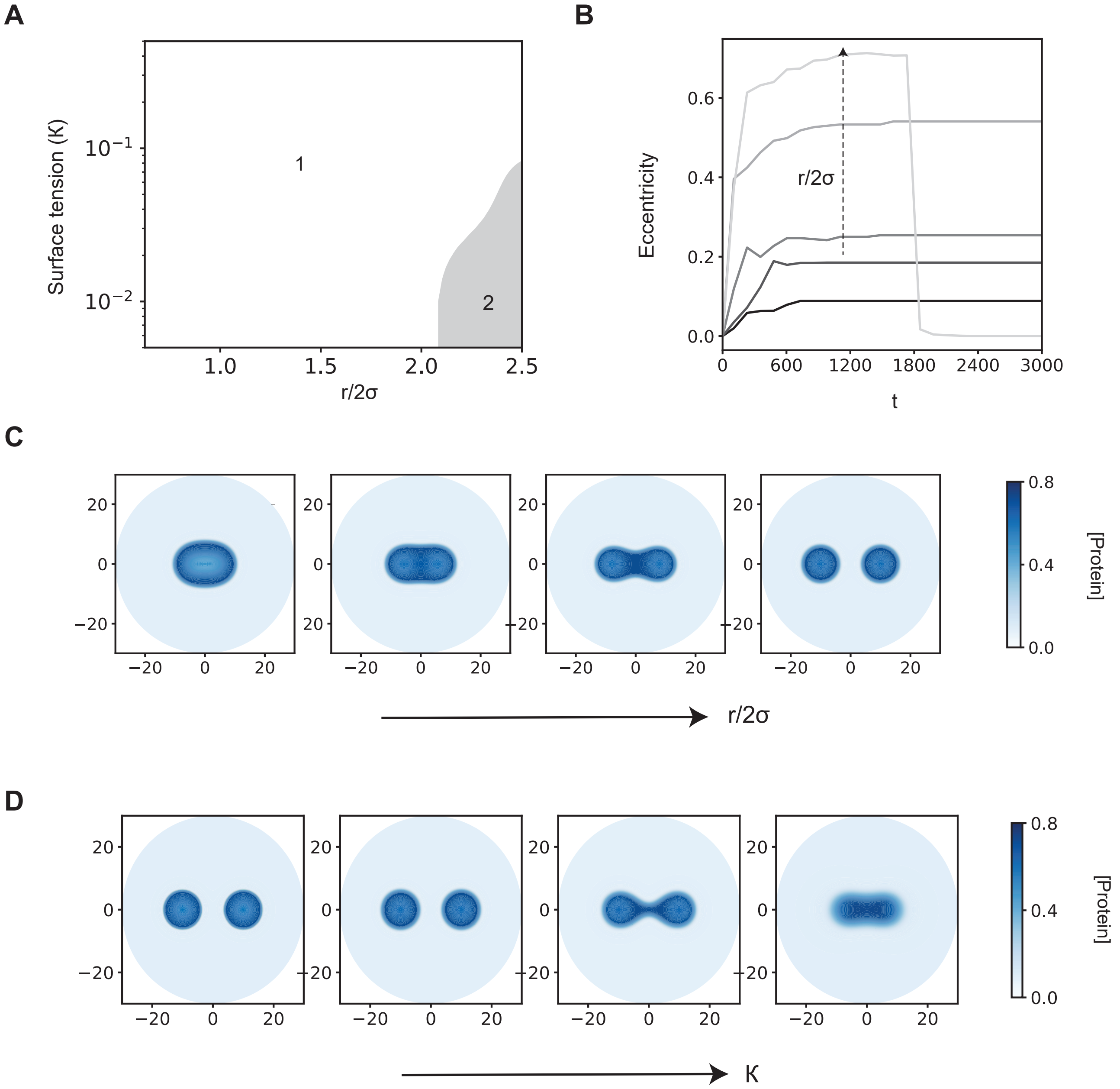


**Figure S4. Two sites of activity can lead to droplet division**

A**.** Phase diagram for condensate morphology upon varying the surface tension of the protein (K) and the distance between two active genomic loci r. The distance between the loci is reported as a dimensionless ratio with the width of the genomic locus (r/2σ). In the parameter regime 1 - we have a single droplet at steady state. In parameter regime 2 where the surface tension is low and the loci are far apart, the system at steady state has two droplets each centered around one of the loci.

B. Dynamics of the eccentricity of the droplets for different distances between the loci (r/2). As the distance between the loci increases, the eccentricity of the droplet increases more rapidly as the droplet reaches a deformed steady state shape. Beyond a threshold distance (light grey), the single droplet deforms so much after a certain time that it quickly splits into two spherical droplets which is a more energetically favorable configuration.

C. Steady state droplet morphology upon varying distance between the loci. Increasing the distance between the loci leads to a gradual deformation of the droplet. Beyond a threshold distance, the droplet splits into two, each one centered around one of the loci.

D. Steady state droplet morphology upon varying the surface tension (). Increasing the surface tension penalizes interface formation and progressively favors a single droplet configuration.
