## Supplementary material for "Organization and regulation of nuclear condensates by gene activity": Unembedded_main_figures

### Figure 1: Model of active nuclear condensates

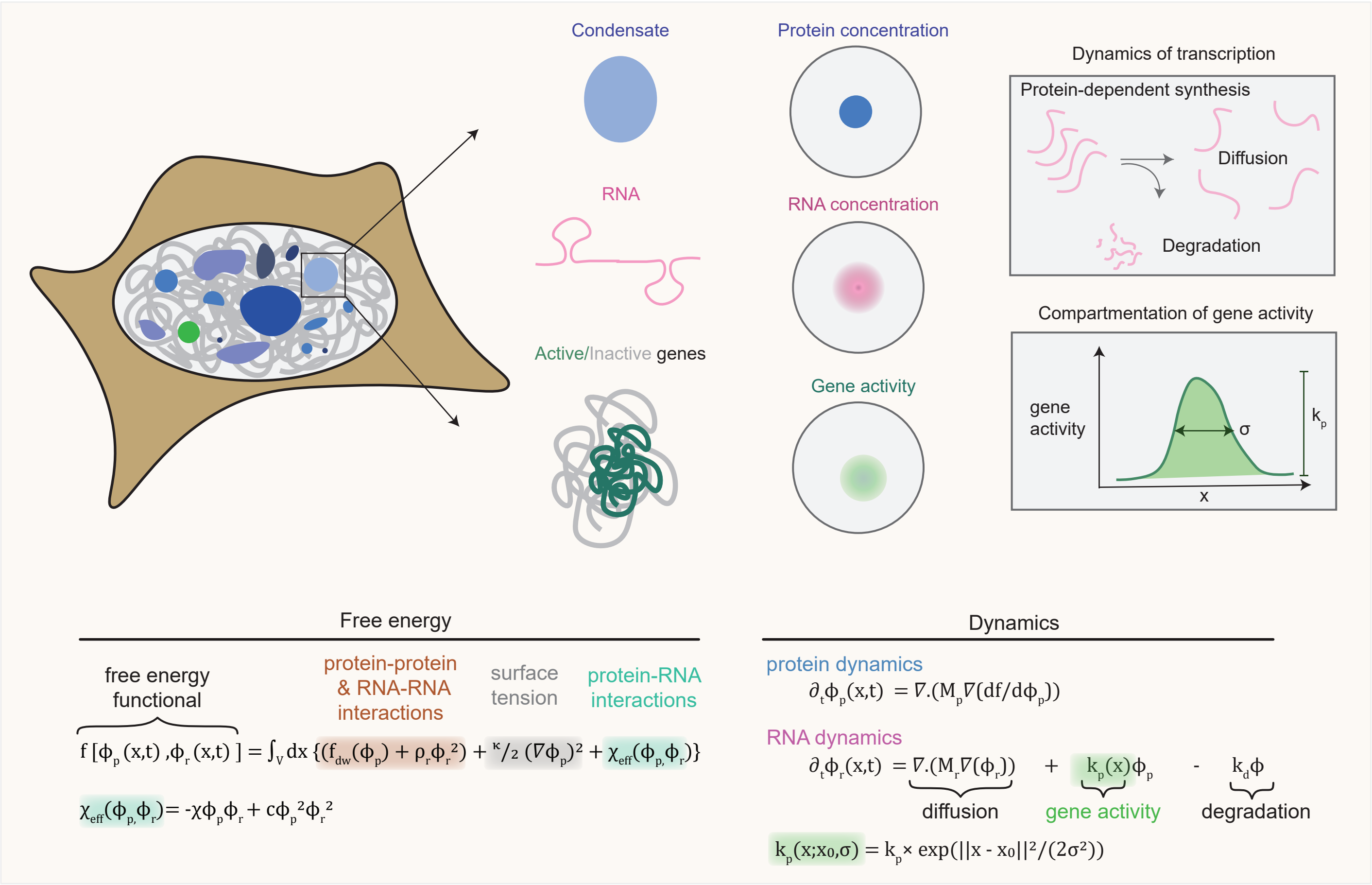

**Figure 2: Spatial clustering of gene activity dictates condensate size and nucleation**

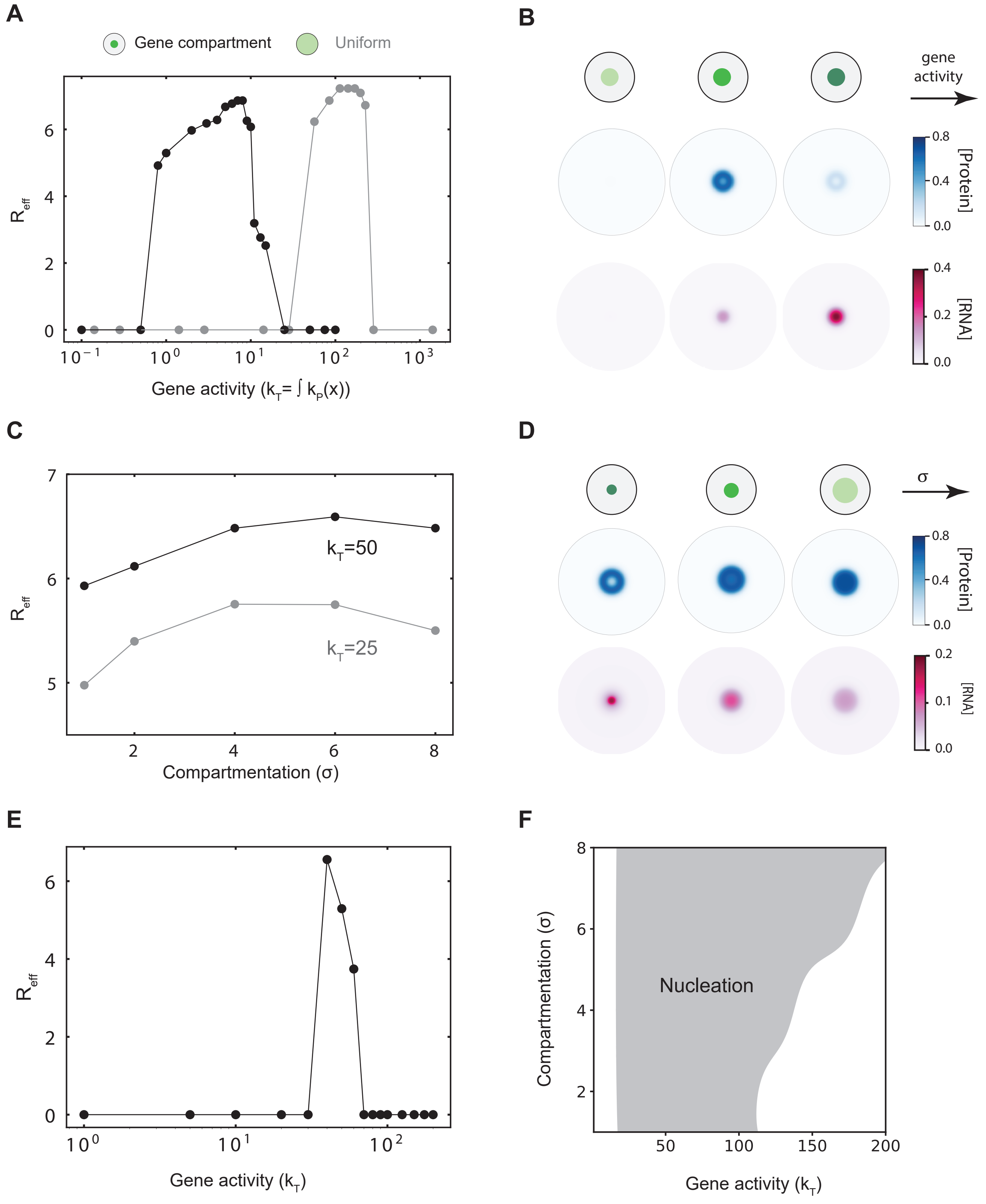

**Figure 3: Active nuclear condensates exhibit unusual steady-state morphologies**

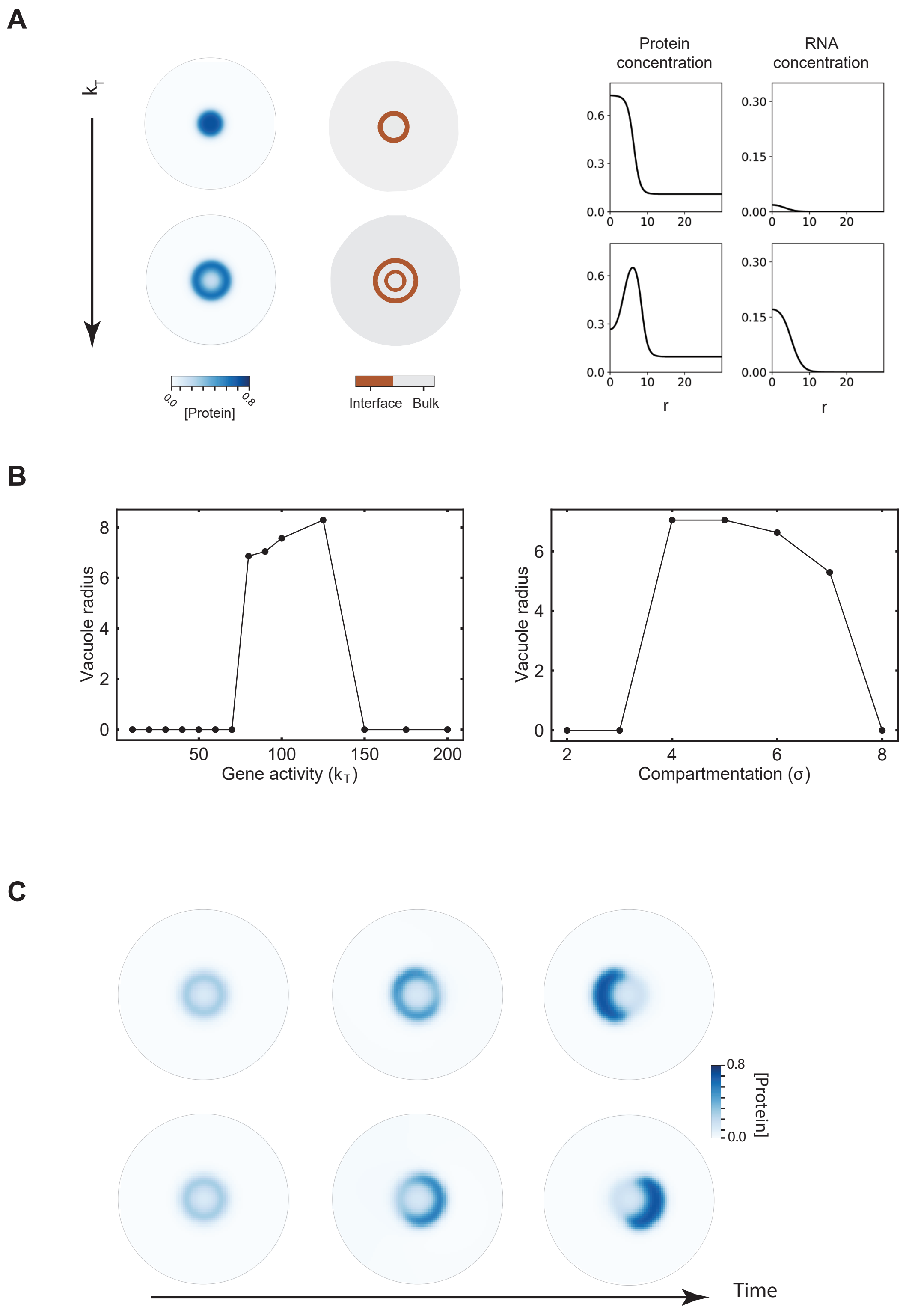

### Figure 4: Distant gene activity induces flow of nuclear condensates

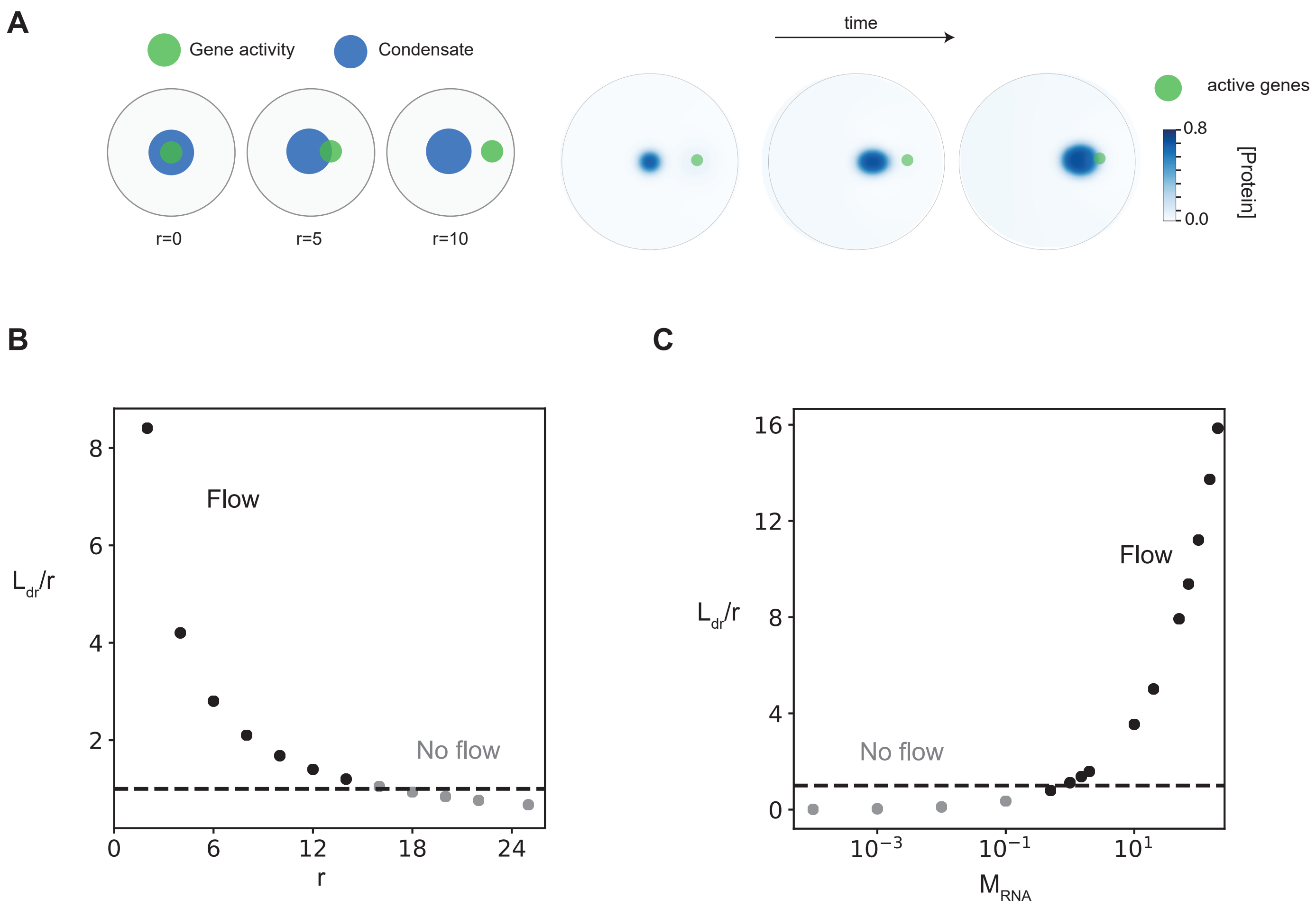

### Figure 5: Gene activity and position dictate emergent condensate morphology and dynamics

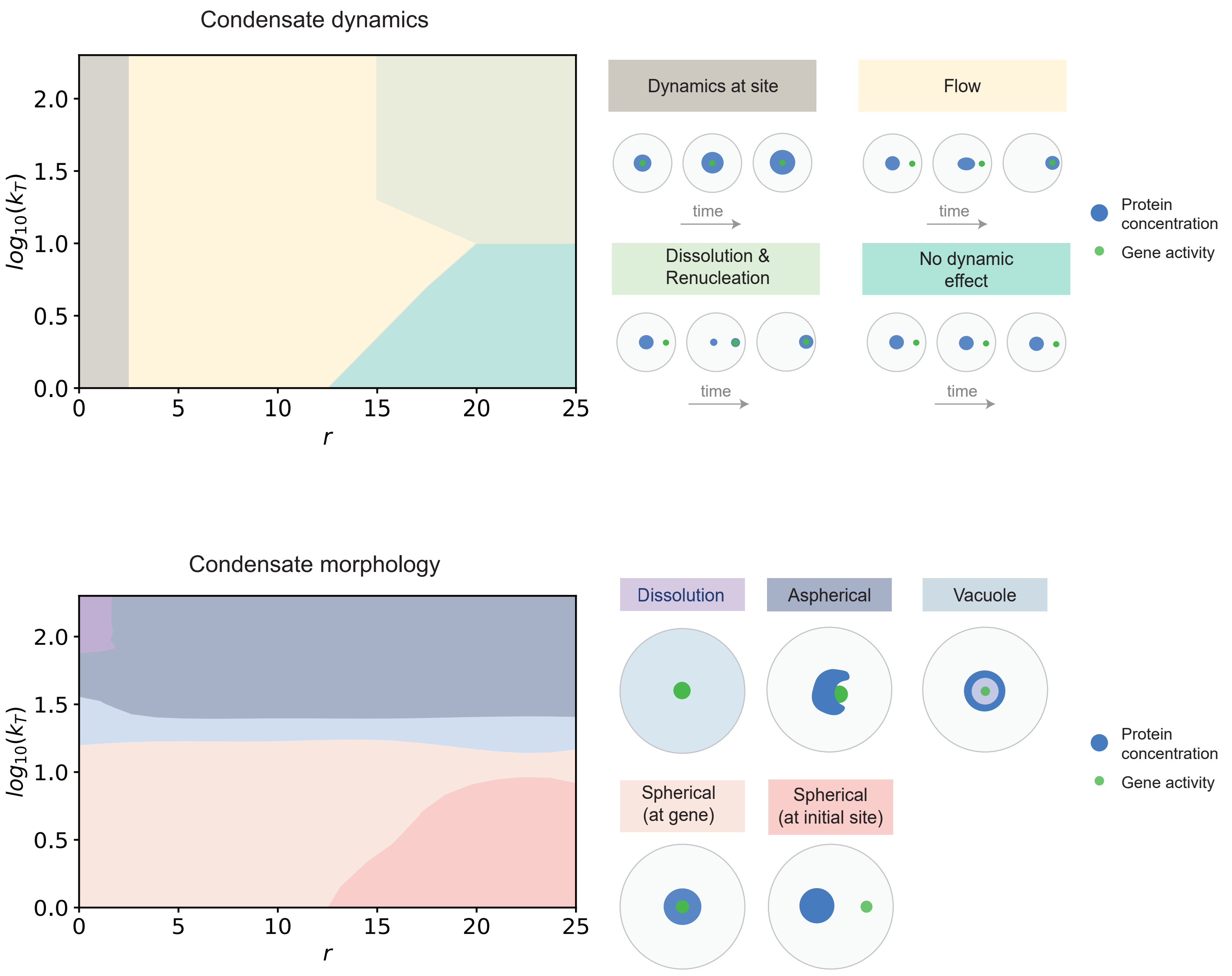

#### Figure 6: Multiple sites of RNA activity compete for or share nuclear condensates

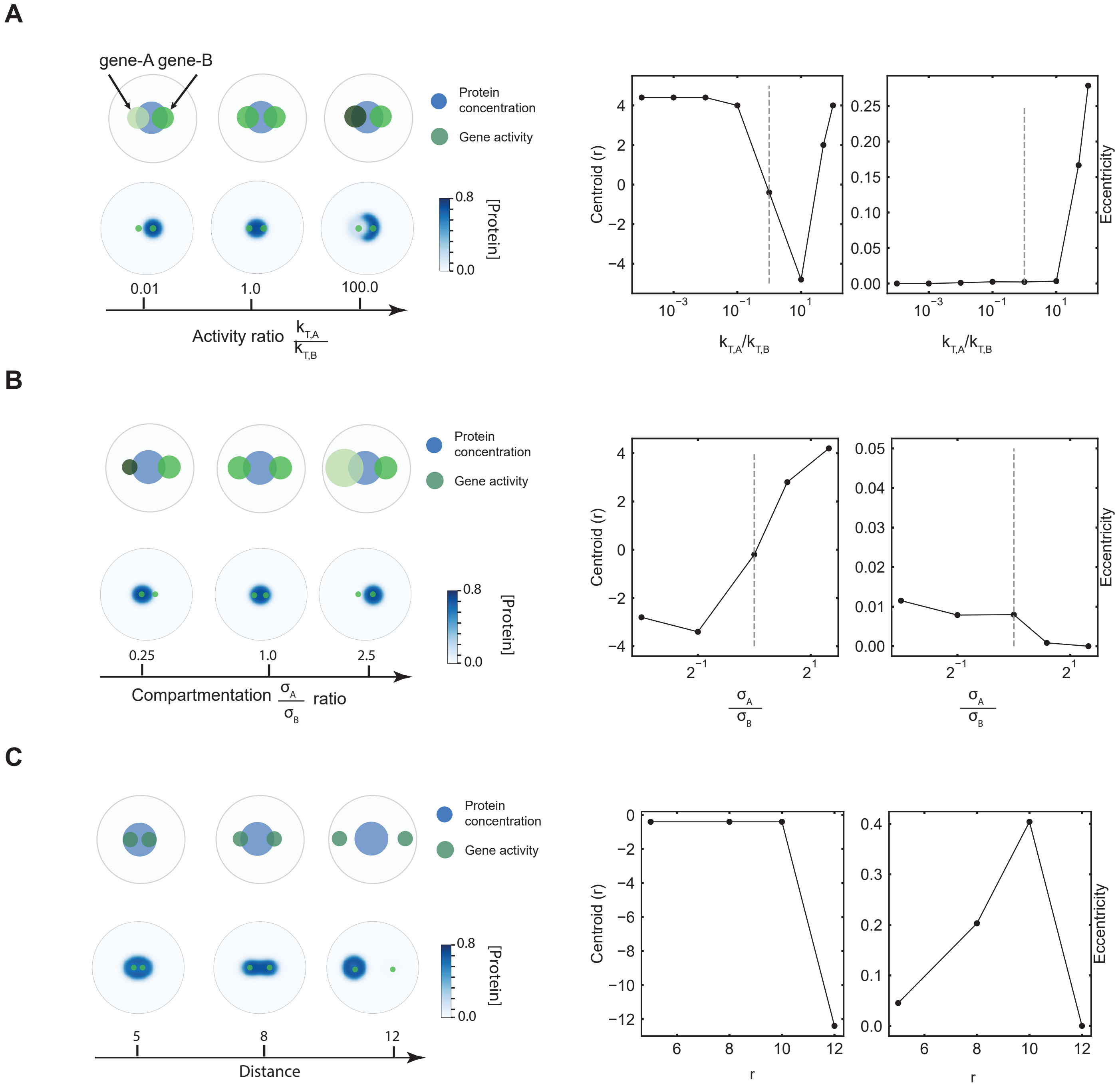
